# Polyvinyl Chloride Degradation by Intestinal *Klebsiella* of Pest larvae

**DOI:** 10.1101/2021.10.03.462898

**Authors:** Zhang Zhe, Haoran Peng, Dongchen Yang, Guoqing Zhang, Jinlin Zhang, Feng Ju

**Affiliations:** Key Laboratory of Coastal Environment and Resources of Zhejiang Province, School of Engineering, Westlake University, 18 Shilongshan Road, Hangzhou 310024, Zhejiang Province, China; Institute of Advanced Technology, Westlake Institute for Advanced Study, 18 Shilongshan Road, Hangzhou 310024, Zhejiang Province, China; College of Plant Protection, Hebei Agricultural University, Baoding 071000, China; Westlake Laboratory of Life Sciences and Biomedicine, Hangzhou, China

**Keywords:** *Spodoptera frugiperda*, intestinal microorganisms, polyvinyl chloride (PVC), *Klebsiella*, degrading enzymes, multiomics

## Abstract

Microbial degradation of polyvinyl chloride (PVC) is eco-friendly and economically attractive but extremely challenging due to the lack of a molecular understanding of the degrading strains and enzymes. Motivated by the serendipitous discovery that the larva of an agricultural invasive insect pest, *Spodoptera frugiperda,* effectively survived PVC film alone, we profiled the intestinal microbiota of *S. frugiperda* larva and screened for PVC-degrading strains. Feeding on PVC film significantly changed the larval intestinal microbiota through selective enrichment of *Enterococcus*, *Ochrobactrum*, *Falsochrobactrum*, *Microbaterium*, *Sphingobacterium* and *Klebsiella*. From the larval intestine, we isolated the biofilm-forming *Klebsiella* sp. EMBL-1 and experimentally verified it as the first *Klebsiella* bacterium known to actively degrade and utilize PVC by various classic physicochemical and morphological analyses. We further used multiomic analyses, complementarily integrating genomic, transcriptomic, proteomic, and metabolomic insights, to identify enzyme-coding genes responsible for PVC degradation and proposed a biodegradation pathway for the bacterial strain. Overall, both *S. frugiperda* and strain EMBL-1 are first found to survive effectively on PVC film by using the polymer as the sole energy source. Moreover, this work exemplifying PVC biodegradation provides a reference for discovering more microbes and enzymatic resources for degrading other recalcitrant plastics.

## Introduction

The globally increasing accumulation of wasted plastics and the associated pollution are serious eco-environmental and socioeconomic problems ^1^. Polyvinyl chloride (PVC) is one of the six widely used plastic polymers, as along with polyethylene (PE), polystyrene (PS), polypropylene (PP), polyurethane (PUR) and polyethylene terephthalate (PET). Among these polymers, the market share of PVC (10.0%) is lower than those of only PE (29.7%) and PP (19.3%) based on the European polymer demand^2^, leading to the generation of tremendous amounts of plastic waste. Landfill and incineration processes are commonly used for the treatment and final disposal of plastic wastes. However, these energy-intensive technologies and industries unfortunately treat plastics solely as useless waste rather than recyclable resources. Worse still, numerous harmful secondary pollutants (e.g., chloride and dioxins) and greenhouse gases are continuously released in large amounts into water, soil, and air environments^3^, making these economically unattractive processes eco-environmentally risky and unsustainable. Unfortunately, there is practically no green or sustainable approach available for disposal of PVC wastes, indicating the need for methodological and technological innovations in the treatment and recycling of such man-made plastics.

Biological treatment and recycling of plastic wastes is a promising approach for the future development of a circular green economy ^4,5^. However, the current research progress on the biodegradation of PVC polymers lags far behind that on the biodegradation of PE^6–9^, PET^10–12^ and PS ^13,14^ Unlike PET, PVC does not have the ester bond which can be hydrolyzed, making its degradation more challenging. Although there is some reported research on the biodegradation of PVC materials, including plasticizers, by microbial consortia^3,15^, little is known about the PVC-degrading microbes or biodegradation pathways involved. Other research has proposed the possibility of PVC biodegradation by several fungal (i.e., Basidiomycotina, Deuteromycota, Ascomycota) or bacterial (i.e., *Pseudomonas*, *Mycobacterium*, *Bacillus* and *Acinetobacter*) taxa based solely on empirically observed morphological and physicochemical changes (e.g., surface damage and molecular weight loss) that visually signify plastic degradation^16–19^. In particular, there is no report on microbial or bacterial genes and enzymes involved in in the degradation of PVC ^20^, excluding Nazia Khatoon’s documentation of the PVC-degrading activity of an extracellular lignin peroxidase of the fungus *Phanerochaete chrysosporium* ^21^. Therefore, global efforts are urgently needed to discover PVC-degrading microbes and enzymatic resources and elucidate the underlying biodegradation mechanisms to develop targeted engineering tools and biotechnologies for PVC waste treatment and recovery.

During laboratory cultivation of the agriculturally invasive insect *Spodoptera frugiperda*, we serendipitously discovered that the pest larva actively bit and fed on PVC film (Figure 1a). Motivated by curiosity and recent reports that some insect species (particularly the larvae of wax moths and meal moths^22^) can consume plastic polymers of PE or PS^13,14,23,24^, we specifically designed triplicate cultivation experiments to check whether *S. frugiperda* larvae could survive solely on PVC film and whether the intestinal microbiota of the larvae may play a role in film digestion, leading to the discovery of the intestinal microbiota-dependent ability of *S. frugiperda* larvae to degrade PVC film for effective survival. Furthermore, using PVC film as the sole energy and organic carbon source, we successfully isolated the first PVC-degrading a *Klebsiella* strain (named EMBL-1) from the intestinal microbiota of the larvae. We explored the biodegradation enzymes and genes and metabolic products of PVC film and proposed the first metabolic pathway for bacterial PVC biodegradation based on multiomic approaches. This work systematically exemplifies the integrated use of universal multiomic approaches for comprehensively mining strains, genes and enzymes with the ability to biodegrade plastics, in addition to providing a theoretical basis for biodegradation-based enzymatic recycling of plastic wastes in the future, as has been recently demonstrated for PET ^13–15^.

**Figure 1.**
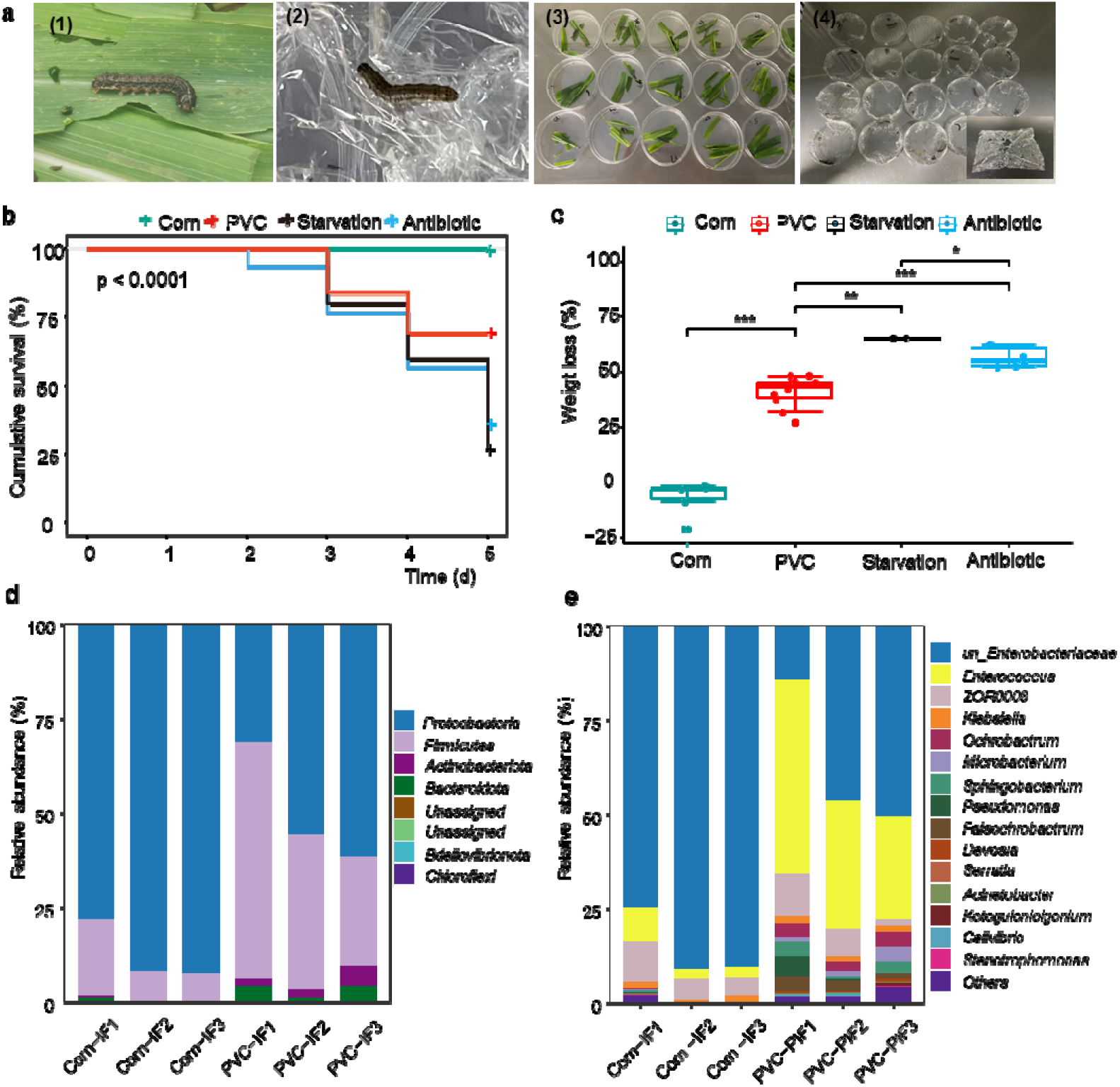
Laboratory cultivation and intestinal microbiota composition of *Spodoptera frugiperda* larvae feeding on corn leaf and PVC film. a, Laboratory feeding of *S. frugiperda* larvae with corn leaves (Corn group) and PVC film (PVC group). b-c, Cumulative survival and body weight loss of larvae in the Corn group, PVC group, Starvation group (no feeding) and Antibiotic group (gentamicin pretreatment of the intestinal microbiota before PVC feeding). d-e, Intestinal microbiota composition at the phylum (d) and genus (e) levels.

## Results

### Discovery and verification of *S. frugiperda* larval survival fueled by PVC film biodegradation

To verify our discovery that larvae of *S. frugiperda* can consume PVC film for effective survival, laboratory cultivation experiments were specifically designed and conducted in triplicate to compare the key physiological indexes (i.e., survival rate, body weight and body length) and intestinal microbiota among larvae under starvation (Starvation group, n = 15), feeding solely on PVC film (PVC group, n = 50), and feeding normally on corn leaves (Corn group, n = 35) (Figure 1a). Overall, the survival rate of the larvae in the PVC group (70%) after 5 days of cultivation was significantly higher than that in the Starvation group (25%) but lower than that in the Corn group (100%, *P* < 0.0001) (Figure 1b). This result was consistent with the significant increase (*P* < 0.0001) in the body weight (Figure 1c) and length (Figure S1a) of the larval groups in the following order: starvation < PVC < corn. This finding indicated that PVC film can provide energy for and sustain the survival of the larva, although the growth efficiency associated with this specialized feeding on PVC film was lower than that with normal feeding on corn leaves, as also morphologically manifested by the contrasting excreted feces (Figure S1b). Moreover, scanning electron microscopy (SEM) analysis of the PVC fragments recovered from excreted feces showed strong surface damage (Figure S1c). These results together verify our discovery that PVC film biodegradation in the gut fuels the survival of *S. frugiperda* larva.

### Interconnections between the intestinal microbiota and PVC biodegradation

The lack of evidence to date for plastic degradation by germ-free invertebrate larvae generally supports the idea that the intestinal microbiota is the key driver of plastic degradation ^22^. Supporting our hypothesis that the gut microbiota is essential for PVC film degradation by *S. frugiperda* larvae, the larval survival rate after 5 days was significantly reduced because of the gentamicin-mediated inhibition of the intestinal microbiota. Accordingly, the body weight (Figure 1c) and body length (Figure S1a) of the PVC-fed larval group treated with gentamicin pretreatment were significantly lower than those of the group without gentamicin pretreatment. The microbial biomass in the larval intestinal microbiota of the Corn group remained stable during the experiment (from 3.90±0.58 to 3.70±0.63×10^6^ CFUs/piece), while the antibiotic-treated group showed an over 99% reduction in microbial biomass. These results suggest the largely dependence of PVC film biodegradation on the larval intestinal microbiota.

PVC film degradation by the intestinal microbiota can lead to the release of transformation products, which we hypothesize should create new ecological niches for microbiome selection through cross-feeding. Consistent with this hypothesis, 16S rRNA gene amplicon sequencing analysis of the intestinal microbiota showed that PVC film degradation triggered a dramatic compositional shift from a Proteobacteria-dominated (87.5±8.0% to 49.5±16.0%) to a Firmicutes-rich (11.9±7.2% to 44.2±17.0%) microbiota (Figure 1d). Further cross-group comparisons at the genus and amplicon sequence variant (ASV) levels showed that compared with normal feeding with corn leaves, PVC feeding and biodegradation dramatically increased the alpha diversity of ASVs in the larval gut microbiota (i.e., the Shannon’s H index increased from 0.7 to 2.0, and the observed species index increased from 30.0 to 70.0) and largely favored the selective enrichment of unclassified *Enterococcus* (4.7±3.7% to 37.0±19.8%), *Ochrobactrum* (0.1±0.2% to 3.4±0.7%), *Falsochrobactrum* (ND to 2.80±1.40%), *Microbacterium* (0.03±0.05% to 2.23±1.55%), *Sphingobacterium* (0.21±0.30% to 2.55±1.54%) and *Klebsiella* (1.4±0.6% to 1.7±0.4%) (Figure 1e), revealing a close interconnection between the pest intestinal microbiota and PVC biodegradation.

### Discovery of strain EBML-1 as the first PVC-degrading *Klebsiella* bacterium

Because the larval intestinal microbiota of *S. frugiperda* is associated with PVC film degradation, we assumed that the larval intestine represents an important reservoir of PVC-degrading strains and promiscuous enzymatic resources. During laboratory screening, a gram-negative strain (Figure S2a), named *Klebsiella* sp. EMBL-1, formed a visible biofilm on the surface of the PVC film after only 10 days of incubation (Figure 2a), causing visible cracks on the surface of the PVC film (Figure 2b), accompanied by a dramatic increase in the strain biomass concentration, i.e., OD_600_ from 0.20 to about 0.60. The cracks formed during initial film degradation could facilitate further plastic degradation ^25^. The strain was further identified as a new *Klebsiella* bacterium (NCBI accession ID: MZ475068) that is most closely related to *Klebsiella variicola* and *Klebsiella pneumoniae*, based on PCR cloning, sequencing, and phylogenetic analyses of the 16S rRNA gene (Figure S2b).

**Figure 2.**
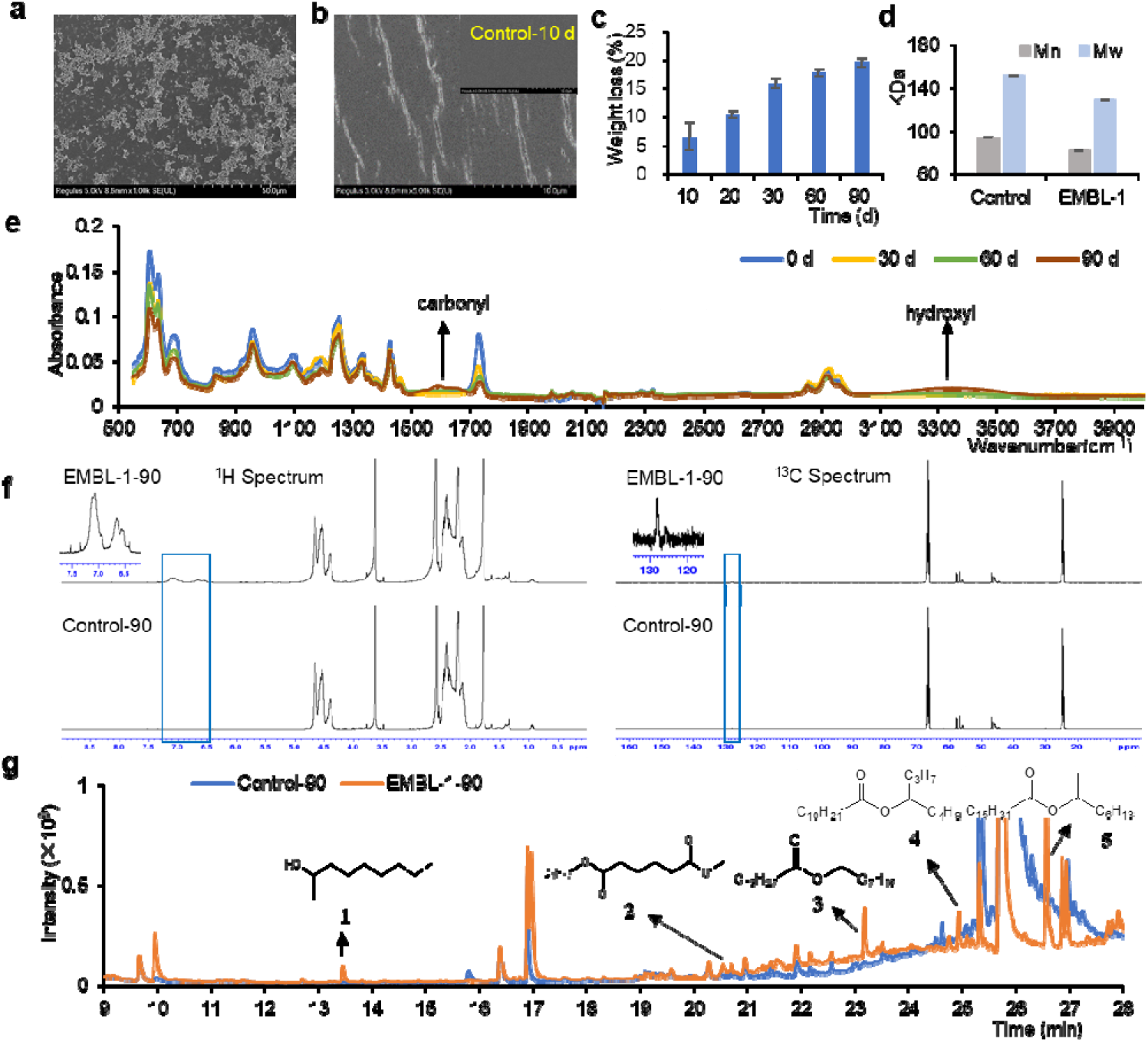
Physicochemical characterization of PVC film degradation by strain EMBL-1. a-b, SEM image of the biofilm formed by EMBL-1 strain on the PVC film on day 10 (a) and SEM image of the cracks formed by EMBL-1 strain on the PVC film on day 10. c, the weight loss curve of PVC film in EMBL-1 group during 90 days. d, the molecular weight of PVC film in the control group and the EMBL-1 group after 90 days. e, the FTIR results for PVC film in the EMBL-1 group over 90 days. f, the results of NMR experiments of PVC in control and EMBL-1 groups after 90 days. g, the detection results of GC-MS of degradation products in PVC films from control and EMBL-1 groups after 90 days.

Using PVC film as a sole energy source, strain EMBL-1 formed a compact biofilm on the film surface after 90 days of incubation. When the biofilm was removed, additional pits and cracks showing film damage were clearly observed (Figure S3a). Based on the contact angle (Figure S3b) and tensile strength tests (Figure S3c) of PVC films, the surface hydrophobicity and tensile strength of the PVC film cultured with strain EMBL-1 changed significantly. These results together suggested that EMBL-1 damaged the physical integrity of the PVC film. During the experiment, the weight loss of the PVC film inoculated with strain EMBL-1 continued to significantly increase over 90 days, and the final average weight loss of the PVC film reached 19.57% (Figure 2c). The results of advanced polymer chromatography (APC, Waters, China) showed that compared with the control group, the molecular weight measures, i.e., Mn and Mw, of the PVC film in the EMBL-1 strain group were decreased by 12.4% and 15.0%, respectively (Figure 2d), indicating that the long-chain structure of PVC was depolymerized, producing lower-molecular-weight fragments. Moreover, thermogravimetric analysis (TGA/DSC 3+/1600 HT, Mettler-Toledo, Switzerland) results showed that the T_max_ and T_onset_ of the PVC film in the EMBL-1 group decreased markedly from 316°C to 279°C and 273°C to 253°C, respectively, while these metrics showed limited changes (T_max_: 316°C to 310°C and T_onset_: 273°C to 265°C) in the control group (Figure S3e-3f), suggesting that the EMBL-1 strain had attacked the PVC polymer chain and reduced the chemical stability of the PVC film.

### Biodegradation products of PVC by strain EMBL-1

Once PVC was proved to be depolymerized by EMBL-1, we further questioned the diversity of the transformation products generated by the strain. Semiquantitative FTIR microspectrometry (Thermo Fisher, Nicolet iS50) analysis showed that the infrared spectrum of the surface of the PVC film inoculated with strain EMBL-1 gradually changed compared with the control group over time (Figure 2e), forming new functional groups such as hydroxyl (3500-3300 cm^−1^) and carbonyl (1550-1650 cm^−1^), indicating microbial oxidation of the PVC film. We then run NMR experiments to determine the structural information of PVC biodegradation products and for the first time made a structure speculation of the corresponding products. On the one hand, the ^1^H NMR and ^13^C NMR spectrum of the pure PVC in EMBL-1 and Control group showed that some substances had been newly produced in the position of 6.5~7.5 ppm (^1^H spectrum) and 128 ppm (^13^C spectrum), suggesting the products have a functional group of -C=C- (Figure 2f). The result of Diffusion Ordered Spectroscopy (DOSY) in the NMR experiments showed the same trend as that of APC, demonstrating not only that molecular weight of PVC film in the EMBL-1 group was lower than that of the Control group, but also that the molecular weight of the newly produced substance was equivalent to the average molecular weight of the EMBL-1 group (Figure S4a). On the other hand, the result of 2D ^1^H-^1^H Correlation SpectroscopY (COSY) indicated the H on the -C=C- was a near-correlated relationship and the structure should be -CH=CH- (Figure S4b). The result of 2D ^1^H-^1^H Nuclear Overhauser Effect SpectroscopY (NOESY) indicated the H on the -CH=CH- functional group had a relationship with the -CH_2_-CH_3_- structure (Figure S4c). Moreover, the results of 2D ^1^H-^13^C Heteronuclear Singular Quantum Correlation (HSQC) spectrum and 2D ^1^H-^13^C Heteronuclear Multiple Bond Correlation (HMBC) spectrum enabled us to determine the relationship between C on the -CH=CH- functional group and C on the -CH_2_-CH_3_- structure, which suggested that the structure of new substance might be –[-CH=CH-CH=CH-]_n_-CH_2_-CH_3_- (Figure S4d-4e). This together with the above results of 2D ^1^H-^1^H NOESY suggested the structure of new substance as –[-CH=CH-CH=CH-]_n_-CH(OH)-CH_2_-, consistent with the generation of hydroxylated products revealed in the FTIR diagram (Figure 2e). The determination of the structure of the new substance also indicated the occurrence of dechlorination reaction. These data altogether also re-verified PVC degradation by EMBL-1.

Further, we used GC-MS to quantitatively profile the degradation products of PVC films over 90 days. By comparative inspection of the peaks with significant differences between the EMBL-1 group and the control group, we identified five potential degradation products (compounds 1 to 5) (Figure 2g), which were sequentially identified as “2-nonanol adipic acid” (1), “adipic acid, methyl octyl ester” (2), “octyl myristate” (3), “dodecanoic acid, isooctyl ester” (4) and “hexadecenoic acid, 1-methylheptyl ester (5)”, respectively, according to the high match score (>800) of each compound in the NIST library (Figure S5). Besides degradation products, three additives, mainly include dioctyl adipate (DOA), dioctyl terephthalate (DOTP) and erucylamide, were also detected from PVC film (Figure S6a-6e and Table S1). Therefore, we set up replicated degradation experiments and found that EMBL-1 had no degradability to the three main additives (Figure S6f and Table S1). In addition, the re-cultivation experiments of EMBL-1 with soybean oil from PVC films showed no cell growth, suggesting that the strain should not derive energy from the residual oil (if any) from the pre-cleaned film (Figure S3d). Considering the fact that other residual impurities have not been detected until now, its influence on the PVC degradation by strain EMBL-1 should be negligible and can be ruled out.

### Genome-level taxonomy and functional profiles of strain EMBL-1

To explore the biodegradation mechanisms of the PVC film, the complete genome of strain EMBL-1 (GenBank Accession CP079802) was constructed based on co-assembly of short reads (150 bp × 2) and long reads (average 27628 bp) derived from Illumina next-generation sequencing and Nanopore third-generation sequencing, respectively. The results showed that the genome contains a 5,662,860 bp circular chromosome with a GC content of 57.31%, and 5646 of the open reading frames (ORFs) were predicted to be protein-coding genes. Interactive tree of life (iTOL) analysis with whole-genome strain information of *Klebsiella* extracted from the Genome Taxonomy Database (GTDB) (Figure 3a) showed that the genome sequence of strain EMBL-1 shared 99.01% average nucleotide identity (ANI) with that of its closest relative, *K. variicola* (RS_GCF_000828055.2).

**Figure 3.**
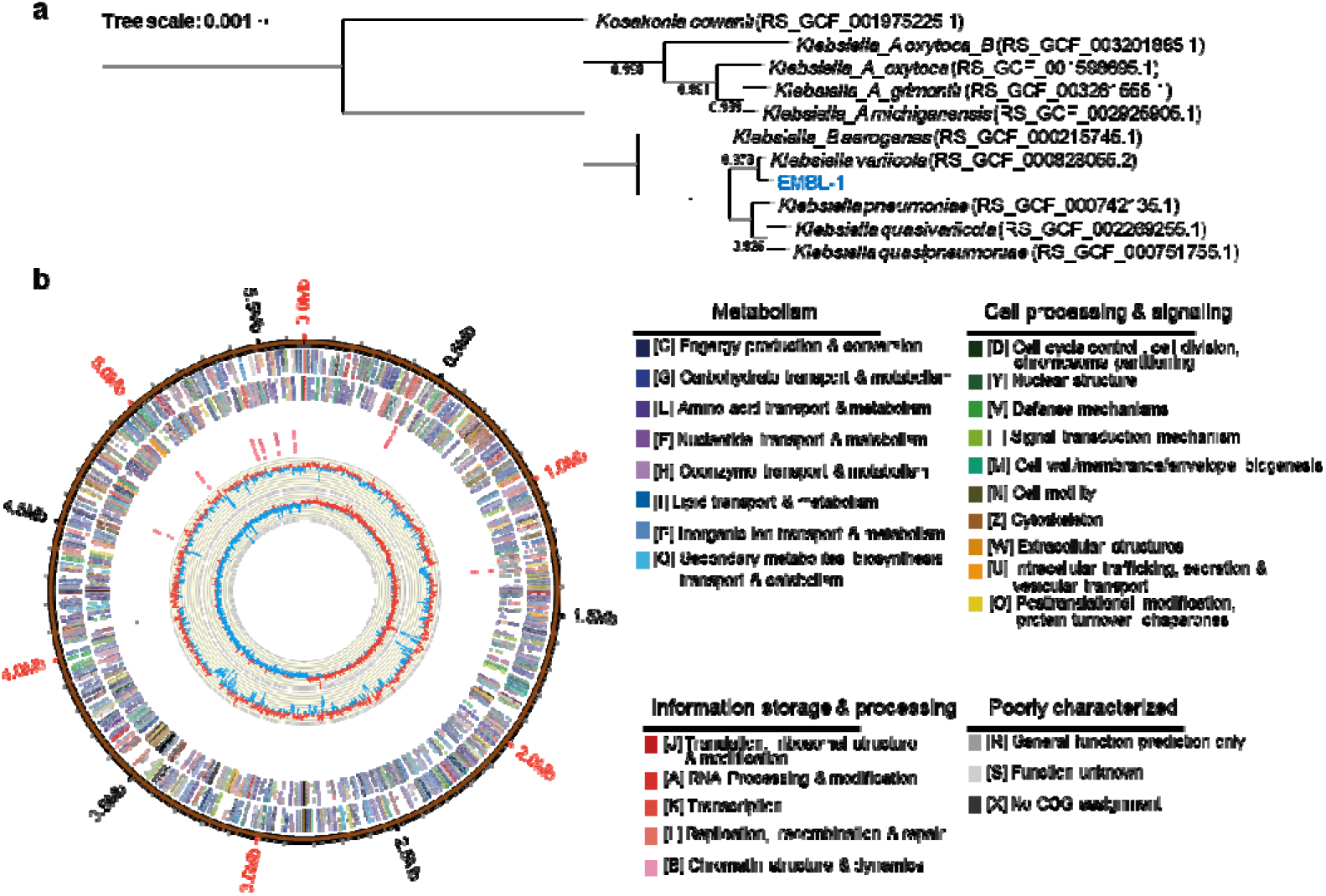
Genome-level phylogeny and functional annotation of strain EMBL-1. a, Phylogenetic tree of the EMBL-1 strain built with iTOL. b, Whole-genome map of strain EMBL-1. Rings from the outside to the center: 1) scale marks of the genome, 2) protein-coding genes on the forward strand, 3) protein-coding genes on the reverse strand, 4) tRNA (black) and rRNA (red) genes on the forward strand, 5) tRNA (black) and rRNA (red) genes on the reverse strand, 6) GC content, 7) GC skew. Protein-coding genes are color coded according to their COG categories.

Functional annotation of the strain EMBL-1 genome (Figure 3b) was performed by homology-based search against four databases, i.e., NR, GO, KEGG and CAZy. Through KEGG metabolic pathway and network analyses, the strain EMBL-1 genome was found to encode 87 genes involved in the metabolism and biodegradation of xenobiotics such as phenylacetic acids and 4-hydroxyphenylacetate. Carbohydrate-active enzyme (CAZy) analysis showed that the genome encoded 74 glycoside hydrolases (GHs), which could contribute to the strong digestion of lignocellulose-containing biomass (e.g., corn leaf) and absorption of carbohydrates by the larvae of *S. frugiperda.* To facilitate the identification of PVC-degrading enzymes, we systematically summarized the reported genera and enzymes related to PE, PVC, and PET degradation and found that strain EMBL-1 had a total of 11 candidate plastic-degrading genes annotated as laccase, alkane monooxygenase, lipase, esterase, peroxidase and carboxylesterase in the KEGG database (Table 1 and Dataset S1), providing first-hand genomic evidence for our finding that the EMBL-1 strain has the ability to degrade PVC polymers.

**Table 1.**
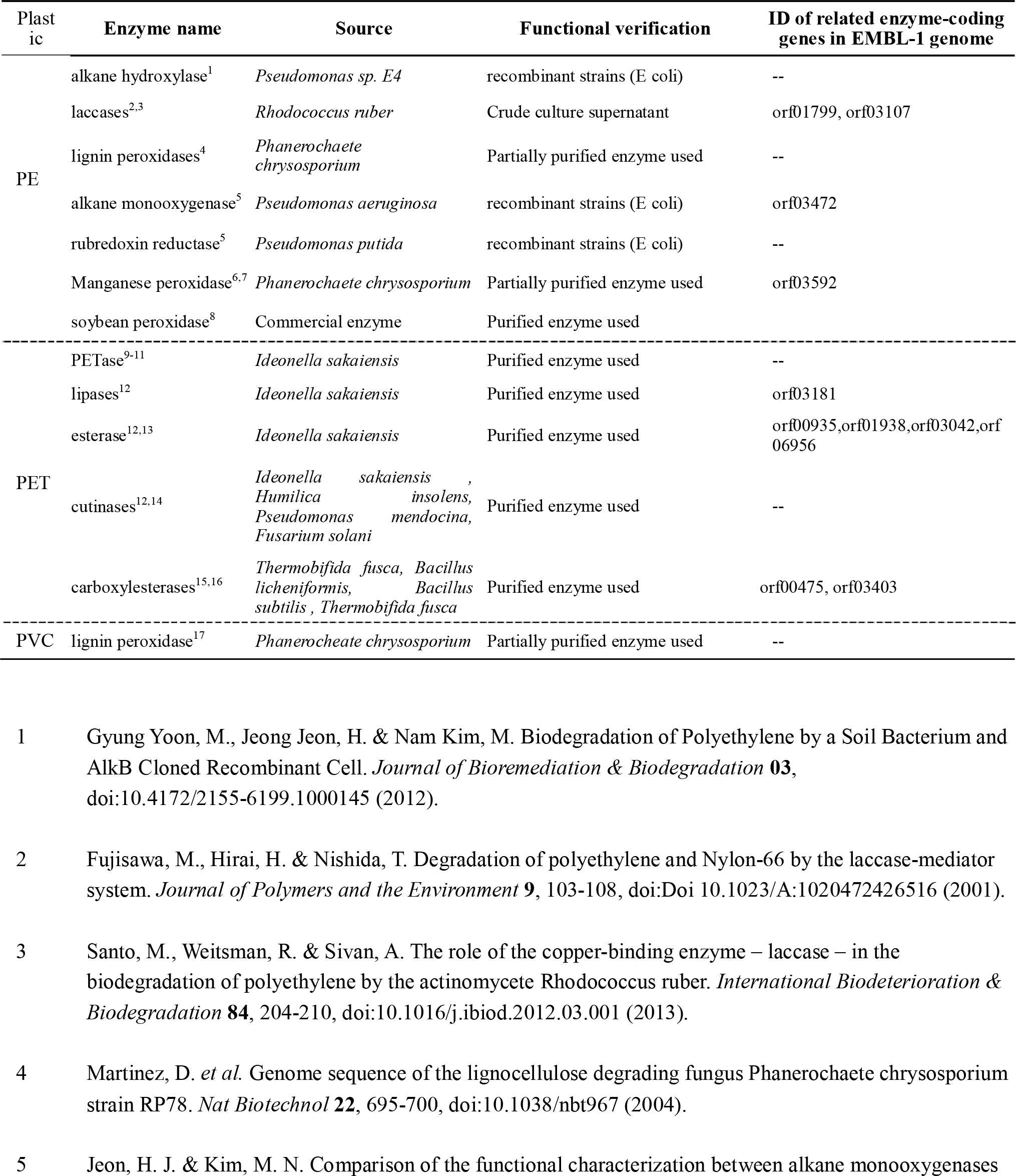
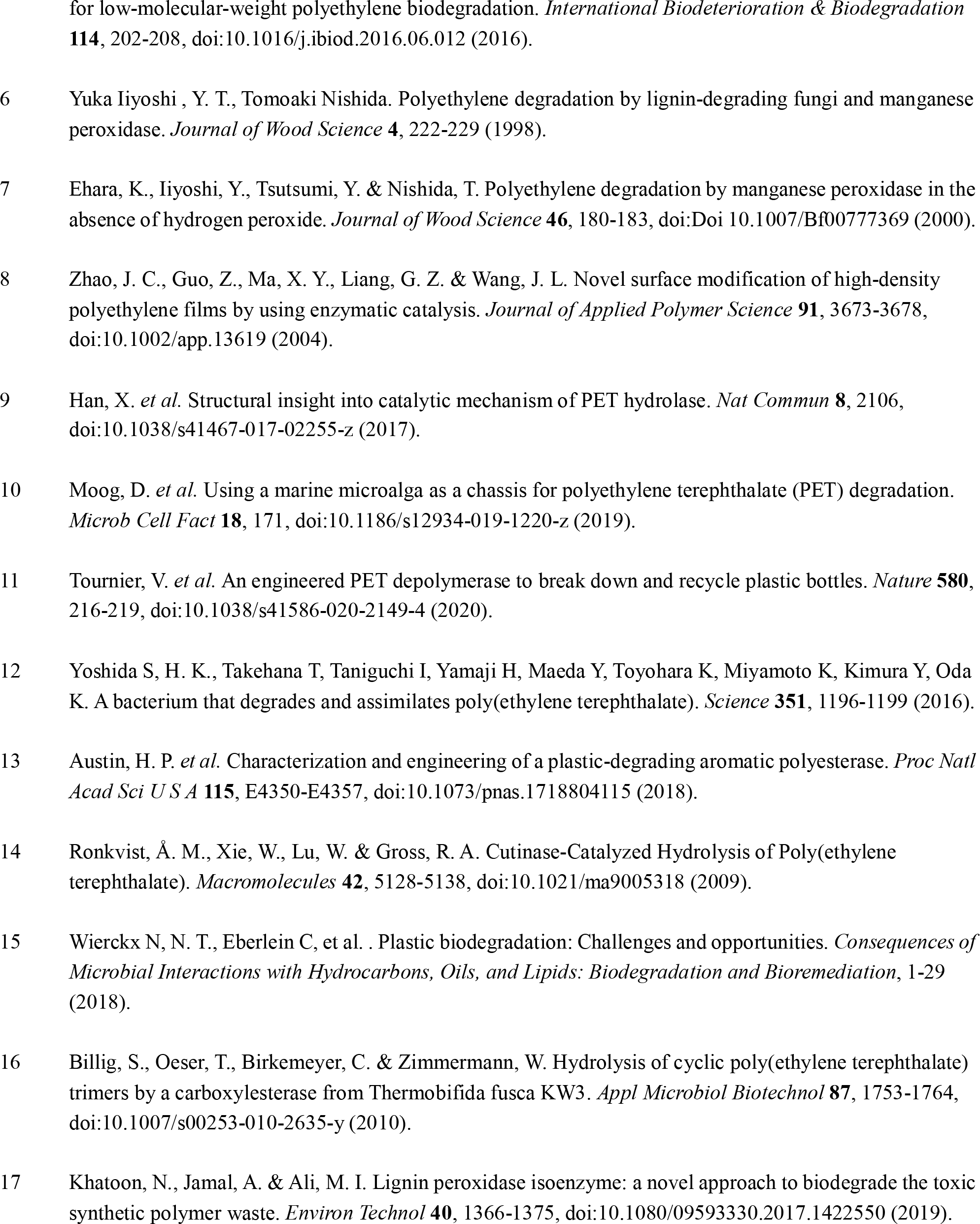
Literature review showing a limited understanding of PVC-degrading microbial strains and enzymes relative to those for PE and PET. Strain EMBL-1 identified in this study is the first experimentally verified PVC-degrading *Klebsiella* isolate. For references of each degrading enzyme, see supporting materials.

### Proteomic analysis of PVC degradation by strain EMBL-1

To further verify the PVC-degrading enzymatic activities, strain EMBL-1 was regrown with PVC film for 30 days before harvesting cells for intracellular (IN) or extracellular (OUT) protein extraction and expression activity tests. The control group supplied with 1% glucose (glu) was set up to better differentiate the metabolic activities for PVC degradation from those for common carbohydrates. The results showed no significant difference in the weight loss of the PVC film between the experimental and control groups (Figure 4a), revealing that an additional organic carbon source did not improve the PVC degradation efficiency of the strain. Instead, glucose addition only increased the glucose metabolism of the EMBL-1 strain (as evidenced by the following proteomic analysis), which in turn increased the protein (a) and biomass (b) levels of the strain (Figure S7). Moreover, the *in vitro* activity measurement showed the PVC-degrading activity of the four protein extracts in the following decreasing order: OUT (13.5%) > OUTglu (10.4%) > IN (5.2%) > INglu (5.0%) (Figure 4b), suggesting that the strain exhibited stronger extracellular than intracellular activities for PVC degradation.

**Figure 4.**
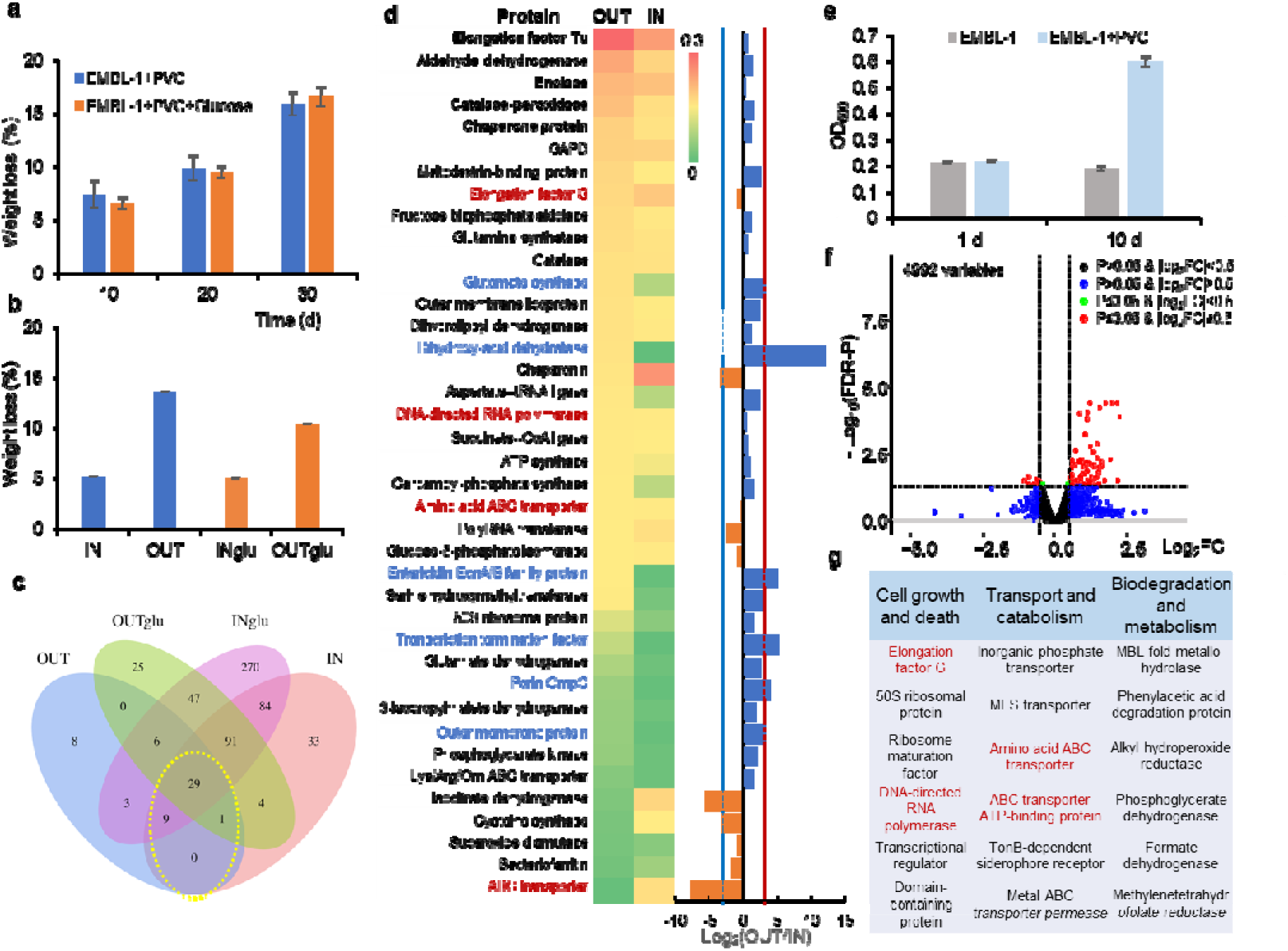
Proteomic and transcriptomic analyses of PVC-degrading metabolic functions of strain EMBL-1. a-b, Weight loss of PVC film under the action of strain EMBL-1 without and with glucose (glu) as supplementary carbon source (a) and four intracellular (IN) or extracellular (OUT) protein extracts under *in vitro* conditions (b). c, Venn diagram showing shared and unique proteins in the protein extracts; d, 39 proteins involved in intracellular and extracellular metabolism of the PVC film. The five most abundant proteins in the OUT extracts are marked in bold, while those that were more upregulated in the OUT extracts than in the IN extracts (OUT/IN≥8) are marked in blue. The proteins marked in red were detected in both proteomes and transcriptomes. e, EMBL-1 cell density (as OD_600_) using PVC film as the sole carbon source. f, Volcano map showing protein-coding genes (red spheres) that were significantly upregulated in the PVC group compared with the control group. g, Functional annotation of upregulated genes, with the genes marked in red being those that were also detected in the proteome.

To comprehensively explore the metabolic functions underlying PVC degradation by strain EMBL-1, we used LC-MS/MS quantitative proteomics to identify a total of 29 proteins jointly expressed in all four experimental and control groups (Figure 4c), plus 10 proteins jointly expressed in only the two PVC-degrading experimental groups (Dataset S2). By inspecting the differential expression of the 39 key proteins between the intracellular (IN) and extracellular (OUT) proteomes, we identified two main categories of proteins associated with PVC degradation. First, we focused on the five most highly expressed extracellular (OUT) proteins (Figure 4d), including i) catalase-peroxidase, enolase, and aldehyde dehydrogenase, which are most likely responsible for the biodegradation of PVC or depolymerized byproducts, and ii) the highly conserved and universal elongation factor Tu and chaperone protein closely related to microbial translation and protective cell responses to nutrient starvation ^27^ and heat shock (or other cooccurring harmful materials such as alcohols, inhibitors of energy metabolism, and heavy metal)^28^, respectively. Among them, catalase-peroxidase has strong redox capacity and polymer depolymerization ability and has also been reported to degrade lignin ^29^, while enolase, with lyase activity, and aldehyde dehydrogenase, with redox activity toward aldehyde groups, are considered putative PVC-degrading proteins. Moreover, we identified another five proteins that were more strongly upregulated (defined here as Log_2_(OUT/IN) ≥ 3) extracellularly (OUT) than intracellularly (IN) (Figure 4d), such as i) dihydroxy-acid dehydratase, which can degrade depolymerized products through cleavage of carbon-oxygen bonds (Figure 5); ii) entericidin EcnA/B family protein, which can manifest the strain’s stress responses to toxic substances (e.g., PVC plasticizers); iii) the porin OmpC and other outer membrane proteins, which can transport some small-molecule metabolites; and iv) the glutamate synthase large subunit, which is well known to be involved in ammonia assimilation ^30^.

**Figure 5.**
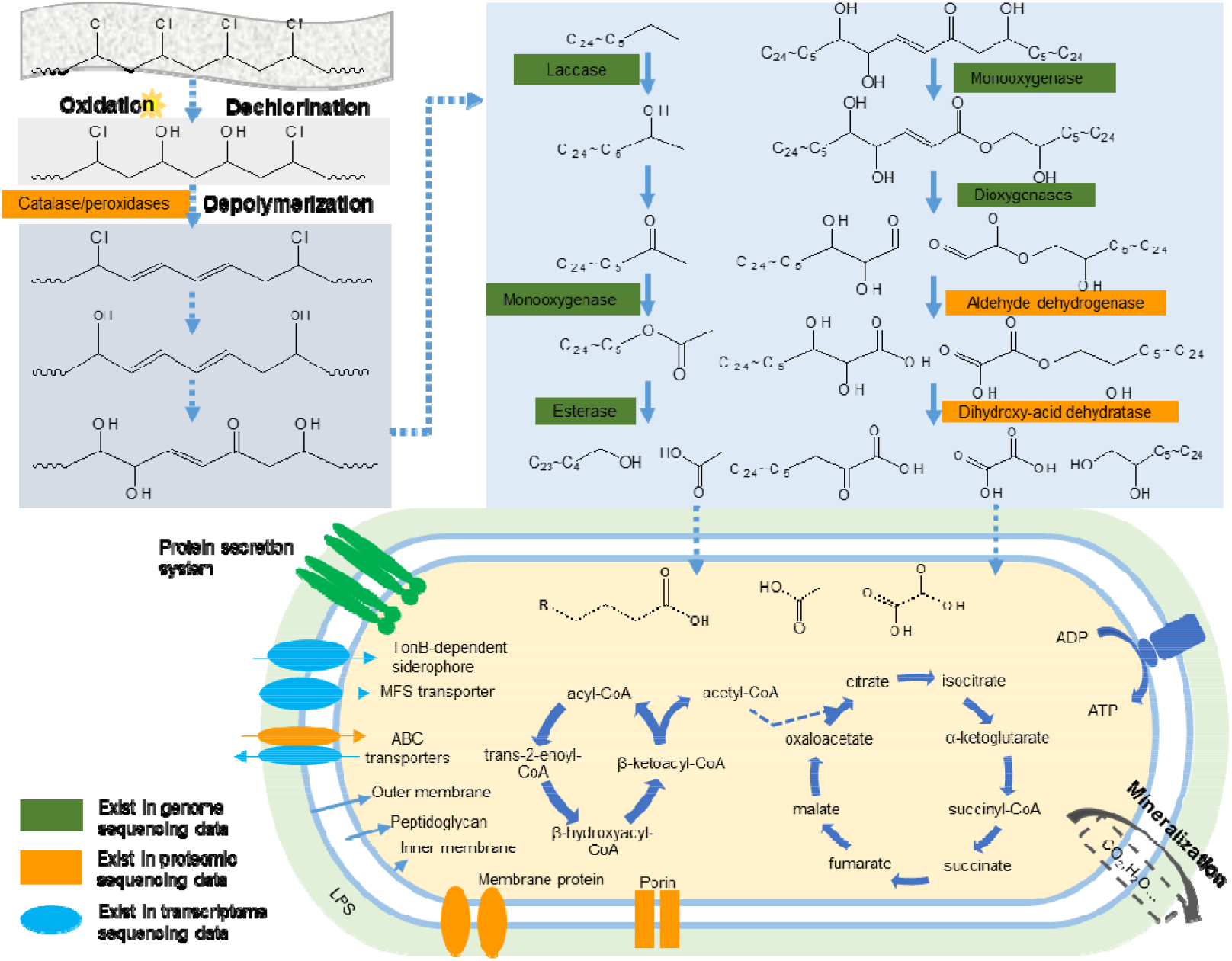
Schematic diagram of the putative PVC degradation pathway of strain EMBL-1. The pathway was proposed based on multiomic analyses that integrated the genomic, proteomic, transcriptomic and metabolomic results for the strain during growth on PVC film.

### Transcriptomic analysis of PVC degradation by strain EMBL-1

To identify enzyme-coding genes associated with PVC film degradation, strain EMBL-1 was first grown on the PVC film in triplicate for 10 days. The weight loss of the film reached approximately 7% (Figure 4a), accounting for an ~3-fold increase in the strain biomass (the OD_600_ increased from 0.20 to 0.58, Figure 4e). We then performed whole-transcriptomic analysis of strain EMBL-1 to screen 77 out of 96 differentially expressed genes that were significantly (FDR-adjusted P ≤ 0.05) upregulated (log_2_(FC) > 0.5) or downregulated (log_2_(FC) < −0.5) in the PVC group (see red spheres, Figure 4f) compared with the control group (Dataset S3). Most of the gene expression changes were ascribed to cell growth and death (e.g., elongation factor G, 50S ribosomal protein, and DNA-directed RNA polymerase), followed by transport and catabolism (e.g., MFS transporter, amino acid ABC transporter, and TonB-dependent siderophore receptor) (Figure 4g). Among them, three transcriptionally active genes were also highly represented in the extracellular (OUT) and/or intracellular (IN) proteomes of the strain (see proteins marked in red, Figure 4d). Notably, biodegradation-related genes, such as MBL fold metallohydrolase, phenylacetic acid degradation protein, and alkyl hydroperoxide reductase, showed active upregulated expression during the strain’s PVC-dependent growth (Figure 4g), revealing their involvement in PVC degradation.

### Multiomic analysis of PVC degradation pathway in strain EMBL-1

Based on the above multiomic approaches that incorporate the complementary results of genomic, transcriptomic, proteomic and metabolite analyses, strain EMBL-1 was found to encode peroxidase, monooxygenase, laccase, lipase, esterase, and carboxylesterase, which are known to degrade PE and/or PET (Table 1 and Dataset S1). These enzymes probably participate in the biodegradation of PVC and its byproducts through a putative pathway that includes responses to abiotic factors, extracellular enzymatic depolymerization, and the intracellular metabolism of degradation byproducts (Figure 5).

First, abiotic factors, including light and oxygen, are widely considered to induce plastic degradation reactions that are initiated via C-C and C-H scission^31,32^. These factors most likely attack and modify the PVC polymer via hydroxylation and carbonylation, as supported by the FTIR. Then, catalase-peroxidase, both known to degrade polymers (e.g., lignin ^29^) and found in our study as the 4^th^ most highly expressed extracellular protein during the PVC-dependent growth of strain EMBL-1 (Figure 4d), depolymerizes the modified polymer into lower molecular weight of polymer (Figure 2d and Figure S4a). After that, oxidation and enzymatic reactions further promoted the degradation of the depolymerized products of PVC, such as the production of -C=C- bonds and hydroxyl groups (Figure 5). The multiomic profiles of other metabolites and degradation enzymes support further stepwise transformation of the long-chain products to shorter products through a series of enzymatic reactions catalyzed by laccase^33^, monooxygenase^34^, dioxygenase, aldehyde dehydrogenase, esterase and dihydroxy-acid dehydratase (Figure 5), such as some C24~C5 byproducts detected by GC-MS/MS analysis (Figure 2g). Significantly, dioxygenase has been reported to modify and degrade plastic polymers via oxidation of C=C functional groups^35^. Although the levels of laccase, monooxygenase, dioxygenase, esterase, and lipase were below the detection limit of the proteomic analysis of EMBL-1, the strain possesses genes encoding these enzymes (Table 1 and Dataset S1). It is likely that the substrate intermediates of these enzymes are promptly consumed by the strain during PVC-dependent growth.

Finally, many genes encoding transport and catabolic proteins were found to be highly represented in the proteomes (Figure 4d) and transcriptomes (Figure 4g) of strain EMBL-1. They are responsible for the transport of small organic molecules and fatty acids to support intracellular catabolism to support strain’s growth.

## Discussion

The limited number of reported PVC-degrading strains originate from natural and nonhost environments such as soil ^19^, landfills ^36^, and marine environments^15,37^, and the underlying biodegradation mechanisms remain unexplored. This study is a new report on the PVC-degrading microbiota and strains, enzymes, and mechanisms in the larval intestines of an agriculturally invasive insect. We discovered that the larvae of *S. frugiperda* could survive solely on the energy derived from PVC degradation by the intestinal microbiota, and further successfully isolated and experimentally verified strain EMBL-1 as the first PVC-degrading *Klebsiella* bacterium. The isolate was taxonomically identified as belonging to *K. variicola* (ANI > 99%). This species is regarded as an emerging pathogen of humans and other animals ^38^. It also harbors plant-associated isolates that can fix nitrogen (to enable its coexistence with plants)^39^ and degrade xenobiotic pollutants (e.g., atrazine ^40^) and natural polymers (e.g., lignin and cellulose ^41,42^). Most importantly, our discovery of strain EMBL-1 and multiomic exploration of its PVC biodegradation mechanisms (as further discussed below) provide a promising methodology, framework and research direction to guide future mechanistic investigations on other PVC-degrading microbes as well as microbial degradation of other plastics.

In addition to the first discovery of the PVC-degrading strain EMBL-1, we also demonstrate an innovative use for multiomic approaches that complementarily integrate DNA, mRNA, protein, and metabolite analyses for mining and elucidating microbial PVC biodegradation genes. The results of our multiomic analyses show that PVC polymers and monomers are effectively utilized by the strain to derive energy for growth. This strain also exhibits a strong adhesion ability and easily forms a biofilm on the PVC film (Figure 2a), which should greatly facilitate its initial destruction of the hydrophobic surface structure of the PVC film by secreting extracellular proteins (as shown by proteomic analysis, Figure 4b–4d), thus allowing its alternative lifestyle, i.e., its ability to utilize PVC for growth^43^. In particular, comparative transcriptomics showed significant upregulation of functional genes involved in xenobiotic biodegradation and metabolism, cell growth and death, and transport and catabolism during cell growth (Figure 4g), revealing that the strain’s strong growth ability and molecular transport ability were closely linked with extracellular PVC polymerization, further conversion, and eventual intracellular utilization.

Until now, except for the sole report on the PVC-degrading activities of fungal lignin peroxidase in *P. chrysosporium* ^21^, little is known about the PVC-degrading pathway, genes and enzymes in bacteria. PVC and PE have similar stable chain structures. Oxygenases, such as monooxygenase and dioxygenase ^44^, are a class of enzymes with relatively high PE degradation activity and can add oxygen to the long carbon chain ^45^, generating free radicals to form carboxyl groups, alcohols, ketones and aldehydes ^26^. The oxidation and cleavage of PE make the polymer more hydrophilic, thus facilitating its contact with other extracellular enzymes (e.g., lipase and esterase) after carboxyl group formation or contact of the amide group with endopeptidase ^26^. In addition, Eyheraguibel *et al.* reported that proteins of the major facilitator superfamily (MFS) or a vector containing an ATP binding cassette (ABC) can integrate low-molecular-weight PE oligomers into cells to achieve PE degradation ^46^. The results of genomic, transcriptomic, and proteomic analyses of the EMBL-1 strain together led us to propose its microbial PVC biodegradation pathway (Figure 5), advancing the current mechanistic understanding of bacterial PVC degradation.

## Conclusion

Microbial biodegradation-based recycling of plastic wastes is promising but extremely challenging, especially when the biodegradation mechanism is unknown. In this study, we started with the discovery of *S. frugiperda* larvae feeding on PVC film, explored the changes in the intestinal microbiota underlying effective larval survival, and, finally, isolated the PVC-degrading *Klebsiella* strain EMBL-1, which exhibited PVC-dependent growth. Through multiomic analyses integrating genome-, transcriptome-, protein- and metabolite-level insights into the strain, we identified a group of functional genes, enzymes, and metabolic pathways closely related to PVC degradation and proposed a hypothetical biodegradation pathway via dechlorination, depolymerization, oxidation and further degradation and mineralization. This study not only paves the way for research into to PVC-degrading microbiota, microbes, and enzymatic resources in the larval intestine of invasive agricultural insects but also provides a multiomic framework and an intriguing scenario that will inspire future studies on the microbial biodegradation of other recalcitrant plastics or xenobiotic contaminants (e.g., pesticides). More importantly, while microbial and enzymatic biodegradation has been demonstrated to be an eco-environmentally sustainable and commercially promising biotechnology for the recycling of PET waste ^10–12^, our study striving to systematically decipher the mechanisms underlying bacterial PVC biodegradation should lay a foundation for follow-up studies to explore and eventually realize sustainable treatment and recycling of PVC plastics enabled by the iterative cycle of “Design-Build-Test-Learn (DBTL)” for DNA, strain and enzymatic engineering.

## Methods

### Field sampling and laboratory cultivation experiments

The larvae of *S. frugiperda* were collected from corn fields in Jiangcheng County, Yunnan Province, China. They were then cultured indoors by feeding on corn leaves (in an artificial insect breeding room with a temperature of 25°C and humidity of 50%-60%). We hypothesized that the intestinal microbiota of *S. frugiperda* larva should play an essential role in digesting PVC film, which enabled the observed and experimentally verified larval survival on PVC film. To test this hypothesis and validate the larval ability to live on PVC film, 130 4_th_-instar larvae with the same growth status were divided into three groups: 1) the control group (starvation, 15 pcs), 2) the Corn group (fed corn leaves, 35 pcs), 3) the PVC group (fed PVC film, 50 pcs), and 4) the Atibiotic group (fed corn leaves with gentamicin for 3 d, and then PVC film, 30 pcs). The weight and body length of all the numbered larvae were measured after 24 h of starvation and after the 5-day experiment. By the end of the experiment, the excreted feces of each experimental group were collected, and the number of survivors was counted. The numbers of culturable cells in the intestinal microbiota in Antibiotic group and Corn group were also determined. Subsequently, the surviving larvae in the Corn group and PVC group were dissected under aseptic conditions to obtain intestinal samples (see details in Method S1), which were labeled and temporarily stored at 4 □ until further operation. The intestinal fecal (IF) samples collected from the PVC and Corn groups were labeled PVC_IF and Corn_IF, respectively. In addition, residual PVC fragments were also recovered from the excreted feces of the PVC group to characterize their surface morphological changes by using a Hitachi field emission scanning electron microscope (Regulus 8230, Japan).

### Exploration between the intestinal microbiota and PVC biodegradation

#### 16S rRNA gene amplicon sequencing

Total DNA was extracted from intestinal fecal samples collected from the experimental groups using a QIAamp Fast DNA Stool Mini Kit following the manufacturer’s recommendations (QIAGEN GmbH, Germany). Then, the hypervariable V4-V5 regions of the prokaryotic 16S rRNA gene were amplified using 515F (5’-GTGYCAGCMGCCGCGGTAA-3’) and 926R (5’-CCGYCAATTYMTTTRAGTTT-3’). The amplicon products of each sample were evenly mixed and sequenced using a paired-end sequencing strategy (PE250) on the Illumina HiSeq2500.

#### Bioinformatic and statistical analyses

For the 16S rRNA gene amplicon data analysis, FastQC (*v*0.11.9) and cutadapt (*v*1.18) were first used to check the quality of the raw data and excise double-ended primers (fastaq files). Dada2 (*v*1.14) was then used to cluster the input sequence with default parameter settings and for further denoising after importing double-ended data through Quantitative Insight into Microbial Ecology (QIIME2-2020.6) and input-format setting parameters. The next step was to select high-quality areas based on FastQC’s report results. Taxonomic classification was conducted using the qiime2 built-in package, and the feature-classifier classify-sklearn machine learning method was used for taxonomic annotation using the SILVA 138 SSU as the reference database. The generated files were imported into R studio version 1.1.414 (R version 4.0.3), and phyloseq (*v*1.32.0) was used for statistical analysis and visualization of the data. In addition, the survival (*v*3.2.7) and survminer packages were used to calculate and draw the survival curve, while ggplot2 (*v*3.3.3) was used to draw box plots of weight and body length. ANOVA was used to check the significance of differences between experimental groups.

### Enrichment, isolation, and identification of the PVC-degrading strain EMBL-1

To isolate plastic-degrading microbes, the intestinal materials of ten larvae were suspended in 10 mL of PBS and vortexed for 5 min. Then, the intestinal mucosa was removed from the mixed solution. The remaining suspension was used as a microbial inoculum and transferred to a 250-mL flask containing 0.1 g of precleaned PVC film and 100 mL of MSM liquid medium. The cleaning procedure of PVC film was described in detail in Method S2. Then, the cells were cultivated on a shaker (150 rpm/min) at 30 □ and transferred every 15 days. After 45 days, the culture medium was first diluted and then spread onto MSM agar medium plates with PVC film as the sole carbon source for cultivation to enrich PVC-degrading strains. The enriched PVC-degrading strain was further subcultured until a pure colony of the isolate was obtained. Depending on whether the isolates were grown in PVC film-amended liquid MSM, the surface changes of the PVC film were inspected by SEM until a PVC-degrading strain (named EMBL-1) was successfully obtained.

To identify the strain, a near full-length 16S rRNA gene sequence was PCR amplified using the universal primers 27F (5’-AGAGTTTGATCCTGGCTCAG-3’) and 1492R (5’-GGTTACCTTGTTACGACTT-3’). The 16S rRNA gene amplicon sequence obtained from Sanger sequencing was deposited in the National Center for Biotechnology Information (NCBI; accession no.: MZ475068) database and annotated using NCBI’s online Basic Local Alignment Search Tool (BLAST) in August 2021.

### Biodegradation of PVC film by strain EMBL-1

Cultivation experiment was conducted to explore the ability of strain EMBL-1 to degrade the cleaned PVC film. Three experimental groups were designed: 1) MSM liquid medium (20 mL) + EMBL-1 strain (OD600=0.2); 2) MSM liquid medium (20 mL) + EMBL-1 strain (OD600=0.2) + PVC film (weighed); and 3) MSM liquid medium (20 mL) + PVC film (weighed). Each experimental group was prepared in 27 replicates and immediately inoculated with the strain. The following operational procedures were repeated every 10 days: i) the PVC films from three replicates of each group were removed; ii) the degradation of the PVC films was examined by SEM, and the films were then weighed; and iii) half of the MSM liquid medium was replaced. The entire experiment lasted for 90 days, during which period, PVC films were collected and stored on days 10 and 90. The posttreatment of the PVC film and the downstream morphological and physicochemical characterization methods are described in detail in Methods S3 and S4, respectively. For the cleaned PVC film, we performed some additional experiment (Method S5-S7) to exclude the EMBL-1’s degrading activity against the main plasticizers and to eliminate the influence of some of the identified substances in the PVC film during the degradation process.

### Characterization of PVC film damage and biodegradation products

#### PVC film damage

To validate and follow PVC film biodegradation by strain EMBL-1, multiple classic physicochemical methods were used together to analyze the temporal changes in the morphological, compositional, and other physiochemical properties over 90 days. First, colonization by the strain was morphologically characterized by SEM after cell fixation. Moreover, the degradation efficiency of the strain was directly measured based on the weight loss of the PVC film on a 10-day basis. Furthermore, changes in the physical properties of the PVC film were detected by contact angle and tensile strength tests (Method S4). The depolymerization of the plastic materials was also recorded by measuring the change in molecular weight. An FTIR microspectrometer (Thermo Fisher, Nicolet iS50, China) was used to analyze and detect the changes in the surface chemical composition and functional groups of the PVC film. Thermogravimetric analysis (TGA/DSC 3+/1600 HT, Mettler-Toledo, Switzerland) was used to compare the initial degradation temperature and the maximum degradation temperature of the PVC film, and the composition, heat stability, and thermal decomposition of the PVC film and the possible intermediate products were examined. The advanced polymer chromatography (APC, Waters, China) was additionally used to determine the molecular weights of different groups of PVC films. A Bruker NEO 600 MHz NMR spectrometer (600.23 MHz for proton frequency) equipped with a TXI probe and a Bruker NEO 500 MHz NMR spectrometer (500.3 MHz for proton frequency) equipped with a BBO Cryoprobe were used to identify some possible the structure of some degradation products.The specific processing steps and testing conditions of the FTIR, TGA, APC and NMR analyses were described in Method S4.

#### Biodegradation products

To obtain more evidence of the biodegradation of PVC film by the EMBL-1 strain, we detected potential biodegradation products of PVC in the PVC films by GC-MS detection. Degraded PVC films weighing 0.03 g were cut into pieces and mixed with 10 mL of tetrahydrofuran, and the mixture was ultrasonicated for 30 min at room temperature. The extract was concentrated to 0.5 ml by drying with nitrogen gas and mixed with 1 mL of n-hexane to obtain some possible products by vortex and ultrasonication for 10 min. The samples were filtered using a 0.22 μm PTFE syringe filter for subsequent steps^47^. The sample was injected at an initial temperature of 40 □ (hold, 4 min), which was progressively increased at 10 °C per minute and held at 280 °C (hold, 5 min). Moreover, the detector conditions, i.e., the transfer line temperature, ion source temperature, ionization mode electron impact and scan time, were maintained at 250°C, 280°C, 70 eV and 0.3 s, respectively.

### Whole-genome sequencing analysis of the PVC-degrading strain EMBL-1

To further explore the mechanism underlying the biodegradation of PVC film by strain EMBL-1 and discover the PVC-degrading genes or enzymes involved in the process, the TIANamp Bacteria DNA Kit was used to extract the genomic DNA of the strain, and the genomic DNA of the EMBL-1 strain was split into two fetches and sequenced using both Illumina next-generation sequencing (PE150) and Oxford Nanopore (PromethION). The experimental procedures, including sample quality testing, library construction, library quality testing, and library sequencing, were performed in accordance with the standard protocol provided by the manufacturers of the sequencer. The bioinformatic analysis included five major steps: raw data quality control, genome assembly, genome component analysis, functional annotation, and genome visualization. In brief, the quality control of raw short reads from Illumina sequencing and raw long reads from Nanopore sequencing were performed in Fastp 0.19.5 and Mecat 2, respectively. Then, the clean short and long reads were coassembled to reconstruct complete genomes using Unicycle (https://github.com/rrwick/Unicycler) to generate complete sequences. The coding sequences (CDSs) were predicted using Glimmer version 3.02 ^48^. Databases such as KEGG, COG, GO, and CAZy were used for functional annotation. In addition, MUMmer software (*v*3.23) was used to compare the target genome with the reference genome to determine the collinearity between the genomes.

### Proteomic analysis of PVC film degradation by strain EMBL-1

To mine enzymatic activities and metabolic pathways related to PVC degradation, biodegradation of PVC film by the EMBL-1 strain was conducted with and without an additional supply of 1% (*w*/*v*) glucose (to distinguish the proteomic signals of the strain from that of PVC film). Then, both intracellular and extracellular proteins were separately extracted from the cells harvested after 30 days based on the acetone precipitation method. The ability of the protein solutions to degrade PVC film was further tested *in vitro* (Method S8). Moreover, proteins were resolved with a Thermo Ultimate 3000 integrated nano-HPLC system that directly interfaced with a Thermo Orbitrap Fusion Lumos mass spectrometer (LC-MS/MS) to explore some related PVC degradation proteins. Details on the experimental setups and procedures, protein extraction methods, protein activity tests, and proteomic analysis are provided in Method S9.

### Transcriptomic analysis of PVC film degradation by strain EMBL-1

#### Experimental design

To mine genes related to PVC degradation, degradation of PVC film by the EMBL-1 strain was conducted for 10 days with a) MSM medium + EMBL-1 (OD600=0.2) and b) MSM medium +EMBL-1 (OD600=0.2) + PVC film (weighed), with three repeats per group. All the treatment groups were cultured in a shaker (30 °C, 150 rpm). By the end of the experiment, the liquid culture was centrifuged at 4 °C to harvest the cells. The total RNA from each treatment group was extracted, and RNA integrity was assessed using the RNA Nano 6000 Assay Kit for the Agilent Bioanalyzer 2100 system (Agilent Technologies). Sequencing libraries were generated using the NEBNext Ultra Directional RNA Library Prep Kit for Illumina (NEB) and sequenced on the Illumina HiSeq 2500 platform. Sequencing was performed at Beijing Novogene Bioinformatics Technology Co., Ltd. The raw transcriptomic data of 6 samples from the two groups were uploaded to the CNGB database (Sub 022027).

#### Bioinformatic analysis

Raw reads were first filtered using fastp to remove the reads that contained 10 low-quality bases (base quality score less than 20) or lengths shorter than 36 bp. Then, the resulting high-quality (HQ) reads were aligned to the *K. variicola* reference genome (*K. variicola* strain FH-1) using hisat2. After alignment, the read counts for each gene were extracted using htseq-count. The gene expression profiles of triplicate transcriptomes in the two groups were compared with PCoA, which inspected one outlier dataset in each group (due to unexpected experimental errors) that was discarded from downstream analysis. Differential expression (DE) at the gene level in our two groups (group a and group b) was evaluated using edgeR version 3.30.3, implemented in R 4.0.3. The p-values presented were adjusted for multiple testing with the procedure of Benjamini and Hochberg to control the type I error rate, and a cutoff of p ≤ 0.05 was used as a threshold to define differential expression. Kraken2 was used to check for contamination in the RNA-seq data.

### Multiomic prediction of degradation pathway of PVC film

To further explore the mechanism underlying the degradation of PVC film by strain EMBL-1, results from multiomic analyses, i.e., genomic, transcriptomic, proteomic and metabolite analyses, were used together to propose a putative pathway for PVC degradation. In brief, the potential plastic-degrading genes encoded in the EMBL-1 genome (Table 1 and Datasets S1) and the metabolites detected by GC-MS were used together to build a putative PVC degradation pathway. Furthermore, 39 proteins jointly expressed during PVC-dependent growth of strain EMBL-1 (Figure 4d and Dataset S2) were aligned against the 96 differentially expressed genes revealed by transcriptomic analysis (Figure 4f and Dataset S3) using NCBI’s BLAST+ 2.9.0 at an e-value cut off of 0.01, generating a list of gene expression and proteomic changes ascribed to the PVC-dependent metabolism of the strain.

## Acknowledgments

This work was supported by Zhejiang Provincial Natural Science Foundation of China (grant no. LR22D010001 to JF) and the National Natural Science Foundation of China (grant no.: 51908467 to JF and 32100091 to ZZ). We would like to thank Dr. Lihan Zhang and Dr. Yubo Wang at Westlake University for discussion and technical assistance and Prof. Ningyi Zhou at Shanghai Jiao Tong University for discussion and helpful suggestions. We also thank Dr. Xingyu Lu, Dr.Xiaohuo Shi, Dr. Yinjuan Chen and Yu Huang from the Instrumentation and Service Center for Molecular Sciences and Dr. Xiuxiu Zhao, Xue Bai and Jinheng Pan from the Biomedical Research Core Facilities at Westlake University for assistance and discussion during the experiment.

