## Supplementary Information for "Polyvinyl Chloride Degradation by Intestinal *Klebsiella* of Pest larvae"

### Supplementary Methods

**Method S1 Dissection method of the larval intestine of *S. frugiperda***

**Method S2 Pre-treatment method of PVC film**

**Method S3 Post-treatment methods of PVC film**

**Method S4 Morphological and physiochemical analysis methods of PVC film**

**Method S5 The degradation activity evaluation of soybean oil degradation in the PVC film by strain EMBL-1**

**Method S6 Identification and content detection of additives in PVC film**

**Method S7 Evaluation of the degradation activity of strain EMBL-1 on the additives in the PVC film**

**Method S8 Proteomic activity of PVC film degradation by strain EMBL-1**

**Method S9 Proteomic analysis with LC-MS/MS**

### Supplementary Tables and Figure Legends

**Table 1 Literature review showing a limited understanding of PVC-degrading microbial strains and enzymes relative to those for PE and PET**

**Table S1 The detection and analysis results of additives in PVC film by GC-MS analysis**

**Figure S1 Changes in body length of *S. frugiperda* larva and characterization of morphological changes of PVC film in feces**

**Figure S2 Screening and isolation of PVC film degrading strains and phylogenetic analysis of 16S rRNA gene sequence**

**Figure S3 The degradation results of strain EMBL-1 on PVC film**

**Figure S4 The spectrum of NMR experiments of PVC**

**Figure S5 Detection results of degradation products of PVC film by GC-MS**

**Figure S6 The detection and identification results of additives in PVC film and the degradation activity of EMBL-1 strains on main additives**

**Figure S7 The growth curve of the EMBL-1 strain and the concentration of extracted protein in the proteomic experiment**

### Supplementary Methods

### Method S1 Dissection method of the larval intestine of *S. frugiperda*

Firstly, the survived larva were placed onto ice to weaken their activity. Then the larvae were sterilized with 70% alcohol for 2 min and washed 3 times with sterile water. Afterwards, the head of the larvae was clamped and removed away from the body, and the intestine of the larvae was gently gripped with tweezers and pulled out and placed into a sterile 1.5 ml centrifuge tube. The intestines of every 10 larva in the Corn group or PVC groups were combined and counted as one replicate sample. Three replicates in each group were used for intestinal microbial DNA extraction and sequencing, and the rest samples were stored in -80℃ for later use.

### Method S2 Pre-treatment method of PVC film

The PVC film purchased from Sinopec Yanshan petrochemical company (11-12 μm film thickness , about 15% of soybean oil content and some additives of unknown content) was cut into 20*20 mm pieces. The PVC pieces were sterilized in 75% ethanol for 20 minutes, washed 3 times with sterile water and air-dried in an ultra-clean workbench. This cleaning process dissolved and washed way the most soybean oil and impurities in the PVC films. Finally, the cleaned PVC pieces were weighed before experiment.

### Method S3 Post-treatment methods of PVC film

**Cell fixation method on PVC film:** Firstly, the PVC film was rinsed properly using sterile water to remove its thick biofilm. Then the cells were collected and sequentially fixed with 2% glutaraldehyde, 25% ethanol, 50% ethanol, 75% ethanol and 100% ethanol.

**Biofilm cleaning method on PVC film:** the PVC film was mixed with a 2% *w*/*v* sodium dodecyl sulfate aqueous solution, shaked for 4 h, and then rinsed with sterile water until the biofilm on the PVC film was completely removed. The cleaned films were weighed after being dried in a drying oven (50°C) for 24 h.

**Calculation method of weight loss of PVC film**: the weight loss (%) was defined as the difference between the weight loss percentage (calculated as 100%*(Initial weight - Final weight) / Initial weight) of the EMBL-1 group and the Control group.

### Method S4 Morphological and physiochemical analysis methods of PVC film

**Detection method of contact angle and tensile strength tests** The contact angle values of PVC film were measured by automatic contact angle measuring instrument (Dataphysics OCA25, Germany). Changes in tensile strength of PVC film were determined on a universal testing machine (TY8000-A, Tianyuan Testing Machine Co. Ltd., Jiangsu, China) equipped with a 200 N cell. The PVC films (2.5cm*0.5cm) in two groups (collected on 90 d) were tested at room temperature (23℃) with a relative humidity of (50±2)% and 25 mm/min, three replicates in each group. All samples were equilibrated to 50% relative humidity for at least 40 h before analysis.

**Detection method of FTIR** The chemical changes of cleaned PVC films were characterized by ATR-FTIR with a scan range of 4000–500 cm-1 using OMNIC software. For each spectrum, 32 scans were taken.

**Detection method of TGA** Dried plastic PVC films of 5 mg were subjected to thermogravimetric analysis using a Perkin Elmer TGA7 thermal analyzer under a nitrogen atmosphere (gas flow: 40 ml/min). The thermograms were recorded from 50 °C to 800 °C at a heating rate of 10°C/min.

**Detection method of APC** Polymer Chromatography (APC) measurements were carried out with three columns XT900-XT450-XT 200 (2.5um, 4.6*150mm) and a RI detector. Tetrahydrofuran (THF, HPLC) was used as mobile phase (0.4 ml/min) after calibration with polystyrene standards of known molecular mass. Non-incubated PVC film was used as references.

**Detection method of NMR**

The NMR experiments were conducted to study the chemical modifications of the polymers by strain EMBL-1. The residual PVC film (50 mg) in control and EMBL-1 groups were dissolved in tetrahydrofuran (THF) solution (15 mL), and then 75mL of methanol was added to precipitate the PVC polymer. The precipitation was obtained by centrifugation and dried. Subsequently, the polymers(precipitation) were resolved in Tetrahydrofuran-d8 (99.5 atom%, Aladdin–Holdings Group, Beijing) to form solutions of approximately 20 mg residue/mL and then transferred to NMR tubes for analysis. All NMR experiments were performed at 25 °C on a Bruker NEO 600 MHz NMR spectrometer (600.23 MHz for proton frequency) equipped with a TXI probe and a Bruker NEO 500 MHz NMR spectrometer (500.3 MHz for proton frequency) equipped with a BBO Cryoprobe. For 1D ^1^H and ^13^C experiments, 64k complex data points were acquired with 16 and 2880 scans, respectively. 2D ^1^H-^1^H Correlation SpectroscopY (COSY) using the “cosygpppqf” pulse sequence were collected with 2 scans × 2048 data points (F2) × 256 increments (F1). 2D ^1^H-^1^H Nuclear Overhauser Effect SpectroscopY (NOESY) using the “noesygpphpp” pulse sequence were collected with 8 scans × 2048 data points (F2) × 256 increments (F1) with the mixing time of 300ms. 2D ^1^H-^13^C Heteronuclear Singular Quantum Correlation (HSQC) using the “hsqcedetgpsisp2.3” pulse sequence were collected with 4 scans × 2048 data points (F2) × 256 increments (F1). 2D ^1^H-^13^C Heteronuclear Multiple Bond Correlation (HMBC) using the “hmbcgplpndqf” pulse sequence were collected with 16 scans × 2048 data points (F2) × 256 increments (F1). Diffusion Ordered Spectroscopy (DOSY) experiments were using the “ledbpgp2s” pulse sequence to measure the self-diffusion coefficient D, with a relaxation delay of 3.0 s and 8 scans in total.16 linear steps from 2% to 95% of gradient strength and a t2 (F2 dimension) of 16k sampling data points were used. The implemented diffusion time big delta and the diffusion gradient length little delta were 80 and 10 ms, respectively.Then, the spectra were analyzed using MestReNova software (version 12.0.0).

### Method S5 The degradation activity evaluation of soybean oil degradation in the PVC film by strain EMBL-1

To check whether strain EMBL-1 degrade the soybean oil washed way from PVC film, commercial soybean oil was purchased and used for further activity detection(Jiangsu Aikon Biopharmaceutical R&D Co.,Ltd). Then the following experiments were set up in triplicates: 1) MSM liquid medium + EMBL-1, 2) MSM liquid medium+ EMBL-1 + soybean oil (2 mg/L), 3) MSM liquid medium + soybean oil (2 mg/L). All treatments were cultured in a shake (150 rpm, 30℃) for 70 days, and the growth of EMBL-1 in each group were measured. After completing the experiment, statistical analysis was performed on the results to determine the degradation activity of the strains on soybean oil (See Fig. S3a for the specific result). EMBL-1 didn’t show a good growth ability in the MSM liquid medium with soybean oil.

### Method S6 Identification and content detection of additives in PVC film

A gas chromatography-mass spectrometry (GC-MS, Trace1300-ISQ7000, ThermoFisher, Singapore) method was established to analyze the composition of additives in the film. The detection conditions are as follows: TD/Py-GC-MS methods were used for quantitation of additives in PVC film using Thermofisher trace1300-ISQ7000. The oven temperature for TD was programmed for 250℃ for 5 min to 350℃ at 20℃/min, and then held at 1min at 350℃. Then the PY temperature was held at 1min at 610℃ and the interface was set at 300 ℃. Sample was injected into the column (TG-5SILMS (30m, 0.25mm, 0.25um) with helium as carrier gas at a constant flow rate of 1ml.min^-1^. The sample was injected at an initial temperature of 50 ℃ (hold for 2 min) which was progressively increased at 10°C per minute and held at 320°C for 3 minutes. Similarly, the detector conditions such as transfer line temperature, ion source temperature, ionization mode electron impact and scan time were maintained at 300°C, 280°C, 70 eV, and 0.3 s respectively. Further, the spectrum attained from the detected compounds at 20-550 Da was compared with GC-MS NIST library of known compounds to identify major additive composition in the PVC film. The test results showed that there were mainly three plasticizers in PVC film, namely dioctyl adipate (DOA), dioctyl terephthalate (DOTP) and erucylamide.

**Quantitative analysis of major additives in PVC film by GC-MS** The content of three major additives in the film were measured by GC-MS referring to the national standard method titled “Consumer product-Plastics-Rapid screening of phthalates” (GB/T 39110-2020). The additive chemicals including dioctyl adipate (DOA, purity≥98%, powder), dioctyl terephthalate (DOTP, purity≥99%, liquid) and erucylamide (purity≥99%, liquid) were purchased at Energy Chemical company. Each sample was injected into the column (same as the above) with helium as carrier gas at a constant flow rate of 1ml.min-1. The sample was injected at an initial temperature of 180 ℃ (hold for 1min) which was progressively increased at 10°C per minute and held at 300°C (hold for 3 min). Moreover, the detector conditions such as transfer line temperature, ion source temperature, ionization mode electron impact and scan time were maintained at 280°C, 250°C, 70 eV, and 0.3 s respectively.

### Method S7 Evaluation of the degradation activity of strain EMBL-1 on the additives in the PVC film

To check whether strain EMBL-1 degrade the additives in the PVC film, the following experiments were set up in triplicates: 1) MSM liquid medium + EMBL-1 (OD600=0.2), 2) MSM liquid medium+ EMBL-1 (OD600=0.2) + DOA/DOTP/erucylamide (100 mg/L, 20 mg/L, 5 mg/L), 3) MSM liquid medium + DOA/DOTP/erucylamide (100 mg/L, 20 mg/L, 5 mg/L) All treatments were cultured in a shake (150 rpm, 30℃) for 30 days, and the growth of EMBL-1 strain in each group were measured every 5 days.

### Method S8 Proteomic activity of PVC film degradation by strain EMBL-1

To explore protein activities involved in the PVC film degradation by strain EMBL-1, the following experiments were set up in nine replicates: 1) MSM liquid medium + EMBL-1 (OD600=0.2) + PVC film (weighed), 2) MSM liquid medium + EMBL-1 (OD600=0.2) + PVC film (weighed) + glucose (1%, *w*/*v*). All treatments were cultured in a shake (150 rpm, 30℃) for 30 days and PVC films were recovered and weighed every 10 days. The pre-treatment and post-treatment methods of PVC film were the same as described in Method S2 and Method S6, respectively. After the experiment, the cell culture solution in the two groups were collected for the next steps.

**Extraction method of intracellular and extracellular proteins** The collected EMBL-1 cells in two groups were lysed by using Branson SFX550 Ultrasonicator, and then the solution was centrifuged to obtain supernatant for further operation. The collected culture solution in two groups were concentrated 10 times for next step. Acetone precipitation method was used to extract intracellular (IN) and extracellular (OUT) proteins. Specific steps are as follows: a mixture of extraction solution and pre-cooled acetone (*v*:*v*=1:1) was stirring for 1 h at 0℃, and then placed on 4℃ overnight. The mixture in each group was concentrated (10000rpm, 4℃) to harvest the proteins from four groups: IN (intracellular protein in group 1)), OUT (extracellular protein in group 1)), INglu (intracellular protein in group 2)), and OUTglu (extracellular protein in group 2)),respectively.

**Test method of PVC-degradation activity of protein extracts** PBS solution was used to dissolve the above protein extracts. The experiments were set up in triplicate as follows: 1) PBS solution + IN (0.1 mg/mL) + PVC film (weighed), 2) PBS solution + OUT (0.1 mg/mL) + PVC film (weighed), 3) PBS solution + Inglu (0.1 mg/mL) + PVC film (weighed), 4) PBS solution + OUTglu (0.1 mg/mL)+PVC film (weighed), and 5) PBS solution + PVC film (weighed). All treatments were cultured in a shake (150 rpm, 30℃) for 48 h before the PVC films were recovered and weighed in each treatment. The weight loss was used to evaluate the PVC-degradation activity of proteins in the four groups: IN, OUT, Inglu and OUTglu.

### Method S9 Proteomic analysis with LC-MS/MS

The SDS-PAGE was used to separate the protein and stained with Coomassie Blue G-250. The gel bands of target were cut into pieces. Sample was digested by trypsin with prior reduction and alkylation in 50 mM ammonium bicarbonate at 37ºC overnight. The digested products were extracted twice with 1% formic acid in 50% acetonitrile aqueous solution and dried to reduce volume by speedvac Vacuum Concentrator.

**LC-MS/MS analysis** The peptides were separated by a 65 min gradient elution at a flow rate 0.300 µL/min with the Thermo Ultimate 3000 integrated nano-HPLC system which is directly interfaced with the Thermo orbitrap fusion lumos mass spectrometer. The analytical column was a home-made fused silica capillary column (75 µm ID, 150 mm length; Upchurch, Oak Harbor, WA) packed with C-18 resin (300 A, 3 µm, Varian, Lexington, MA). Mobile phase A consisted of 0.1% formic acid, and mobile phase B consisted of 80% acetonitrile and 0.1% formic acid. The mass spectrometer was operated in the data-dependent acquisition mode using the Xcalibur 4.1 software and there is a single full-scan mass spectrum in the Orbitrap (300-1800 m/z, 60,000 resolution) followed by 20 data-dependent MS/MS scans at 30% normalized collision energy. Each mass spectrum was analyzed using the Peak studio for the database searching. The reference strain is *Klebsiella variicola* (strain 118)

**Identification of differentially expressed proteins** Based on the PVC-degradation activity of four proteins (OUT > OUTglu > IN > INglu), statistical analysis was focused on the proteins shared by the OUT group and the IN group which showed degrading activities of PVC film. In total, 39 proteins were firstly selected within the yellow dotted line. Further, the relative abundance of these proteins in their respective groups was calculated and visualized in a heat map. Moreover, the differential profiles in protein expression between the two groups were calculated based on the relative abundance to further narrow the list of potential PVC-degrading proteins (Log_2_(OUT/IN) ≥3).

### Supplementary Tables and Figures

### Table 1 Literature review showing a limited understanding of PVC-degrading microbial strains and enzymes relative to those for PE and PET. Strain EMBL-1 identified in this study is the first experimentally verified PVC-degrading *Klebsiella* isolate.

|  | **Enzyme name** | **Source** | **Functional validation** | **Protein-coding gene ID in the genome of strain EMBL-1** |
| --- | --- | --- | --- | --- |
| PE | alkane hydroxylase^1^ | *Pseudomonas sp. E4* | Recombinant strains (*E. coli*) | -- |
|  | laccase^2,3^ | *Rhodococcus ruber* | Crude culture supernatant | orf01799, orf03107 |
|  | lignin peroxidase^4^ | *Phanerochaete*  *chrysosporium* | Partially purified enzyme used | -- |
|  | alkane monooxygenase^5^ | *Pseudomonas aeruginosa* | Recombinant strains (*E. coli*) | orf03472 |
|  | rubredoxin reductase^5^ | *Pseudomonas putida* | Recombinant strains (*E. coli*) | -- |
|  | manganese peroxidase^6,7^ | *Phanerochaete chrysosporium* | Partially purified enzyme used | orf03592 |
|  | soybean peroxidase^8^ | Commercial enzyme | Purified enzyme used |  |
| PET | PETase^9-11^ | *Ideonella sakaiensis* | Purified enzyme used | -- |
|  | lipases^12^ | *Ideonella sakaiensis* | Purified enzyme used | orf03181 |
|  | esterase^12,13^ | *Ideonella sakaiensis* | Purified enzyme used | orf00935,orf01938,orf03042,orf06956 |
|  | cutinases^12,14^ | *Ideonella sakaiensis, Humilica insolens, Pseudomonas mendocina, Fusarium solani* | Purified enzyme used | -- |
|  | carboxylesterases^15,16^ | *Thermobifida fusca, Bacillus licheniformis, Bacillus subtilis, Thermobifida fusca* | Purified enzyme used | orf00475, orf03403 |
| PVC | lignin peroxidase^17^ | *Phanerocheate chrysosporium* | Partially purified enzyme used | -- |

### Table S1 The detection and analysis results of additives in PVC film by GC-MS analysis

| Name | RT/min | Qualitative and quantitative transition | Standard curve/R^2^ | Content in PVC film (%) |
| --- | --- | --- | --- | --- |
| DOA | 11.78 | 129,57,112/129 | y=2E+06x-226835, R^2^=0.9925 | 22.90 |
| DOTP | 14.53 | 70,112,149/70 | y=952890x-2E+06, R^2^=0.9935 | 5.23 |
| Erucylamide | 15.09 | 59,72,55/59 | y=523400x-1E+06, R^2^=0.9936 | 2.05 |

1	Gyung Yoon, M., Jeong Jeon, H. & Nam Kim, M. Biodegradation of Polyethylene by a Soil Bacterium and AlkB Cloned Recombinant Cell. *Journal of Bioremediation & Biodegradation* **03**, doi:10.4172/2155-6199.1000145 (2012).

2	Fujisawa, M., Hirai, H. & Nishida, T. Degradation of polyethylene and Nylon-66 by the laccase-mediator system. *Journal of Polymers and the Environment* **9**, 103-108, doi:Doi 10.1023/A:1020472426516 (2001).

3	Santo, M., Weitsman, R. & Sivan, A. The role of the copper-binding enzyme – laccase – in the biodegradation of polyethylene by the actinomycete Rhodococcus ruber. *International Biodeterioration & Biodegradation* **84**, 204-210, doi:10.1016/j.ibiod.2012.03.001 (2013).

4	Martinez, D. *et al.* Genome sequence of the lignocellulose degrading fungus Phanerochaete chrysosporium strain RP78. *Nat Biotechnol* **22**, 695-700, doi:10.1038/nbt967 (2004).

5	Jeon, H. J. & Kim, M. N. Comparison of the functional characterization between alkane monooxygenases for low-molecular-weight polyethylene biodegradation. *International Biodeterioration & Biodegradation* **114**, 202-208, doi:10.1016/j.ibiod.2016.06.012 (2016).

6	Yuka Iiyoshi , Y. T., Tomoaki Nishida. Polyethylene degradation by lignin-degrading fungi and manganese peroxidase. *Journal of Wood Science* **4**, 222-229 (1998).

7	Ehara, K., Iiyoshi, Y., Tsutsumi, Y. & Nishida, T. Polyethylene degradation by manganese peroxidase in the absence of hydrogen peroxide. *Journal of Wood Science* **46**, 180-183, doi:Doi 10.1007/Bf00777369 (2000).

8	Zhao, J. C., Guo, Z., Ma, X. Y., Liang, G. Z. & Wang, J. L. Novel surface modification of high-density polyethylene films by using enzymatic catalysis. *Journal of Applied Polymer Science* **91**, 3673-3678, doi:10.1002/app.13619 (2004).

9	Han, X. *et al.* Structural insight into catalytic mechanism of PET hydrolase. *Nat Commun* **8**, 2106, doi:10.1038/s41467-017-02255-z (2017).

10	Moog, D. *et al.* Using a marine microalga as a chassis for polyethylene terephthalate (PET) degradation. *Microb Cell Fact* **18**, 171, doi:10.1186/s12934-019-1220-z (2019).

11	Tournier, V. *et al.* An engineered PET depolymerase to break down and recycle plastic bottles. *Nature* **580**, 216-219, doi:10.1038/s41586-020-2149-4 (2020).

12	Yoshida S, H. K., Takehana T, Taniguchi I, Yamaji H, Maeda Y, Toyohara K, Miyamoto K, Kimura Y, Oda K. A bacterium that degrades and assimilates poly(ethylene terephthalate). *Science* **351**, 1196-1199 (2016).

13	Austin, H. P. *et al.* Characterization and engineering of a plastic-degrading aromatic polyesterase. *Proc Natl Acad Sci U S A* **115**, E4350-E4357, doi:10.1073/pnas.1718804115 (2018).

14	Ronkvist, Å. M., Xie, W., Lu, W. & Gross, R. A. Cutinase-Catalyzed Hydrolysis of Poly(ethylene terephthalate). *Macromolecules* **42**, 5128-5138, doi:10.1021/ma9005318 (2009).

15	Wierckx N, N. T., Eberlein C, et al. . Plastic biodegradation: Challenges and opportunities. *Consequences of Microbial Interactions with Hydrocarbons, Oils, and Lipids: Biodegradation and Bioremediation*, 1-29 (2018).

16	Billig, S., Oeser, T., Birkemeyer, C. & Zimmermann, W. Hydrolysis of cyclic poly(ethylene terephthalate) trimers by a carboxylesterase from Thermobifida fusca KW3. *Appl Microbiol Biotechnol* **87**, 1753-1764, doi:10.1007/s00253-010-2635-y (2010).

17	Khatoon, N., Jamal, A. & Ali, M. I. Lignin peroxidase isoenzyme: a novel approach to biodegrade the toxic synthetic polymer waste. *Environ Technol* **40**, 1366-1375, doi:10.1080/09593330.2017.1422550 (2019).

### Figure S1 Changes in body length of *S. frugiperda* larva and characterization of morphological changes of PVC film in feces.

a, changes in body length of *Spodoptera frugiperda* larvae. b, feces excreted by larva in Corn group (1) and PVC group (2). c, characterization results of degradation PVC film recovered from feces in PVC group, (1)-(4) represented the degradation of PVC film from low magnification to high magnification in SEM.

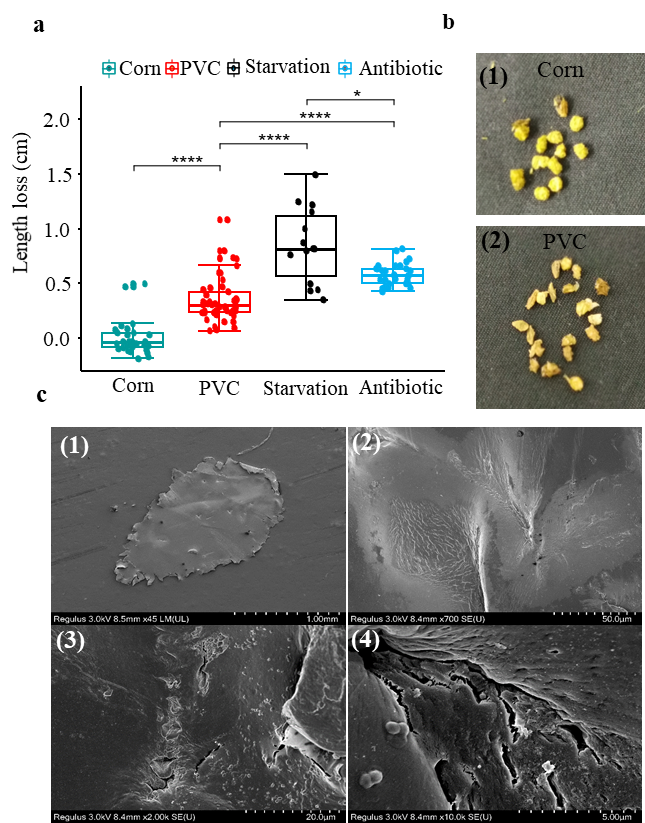

### Figure S2 Screening and isolation of PVC film degrading strains and phylogenetic analysis of 16S rRNA gene sequence

a, experimental screening and isolation. The dissected intestine of larva in PVC group was used to inoculate and enrich for degrading strain EMBL-1, which was cultured on PVC film in MSM liquid medium and LB solid medium. b, phylogenetic tree of 16S rRNA gene sequences showing PVC-degrading strain EMBL-1 as a novel Klebsiella strain most closely related to *Klebsiella variicola* and *Klebsiella pneumoniae*. The analysis was conducted using the Fast Minimum Evolution method at a maximum sequence difference of 0.75.

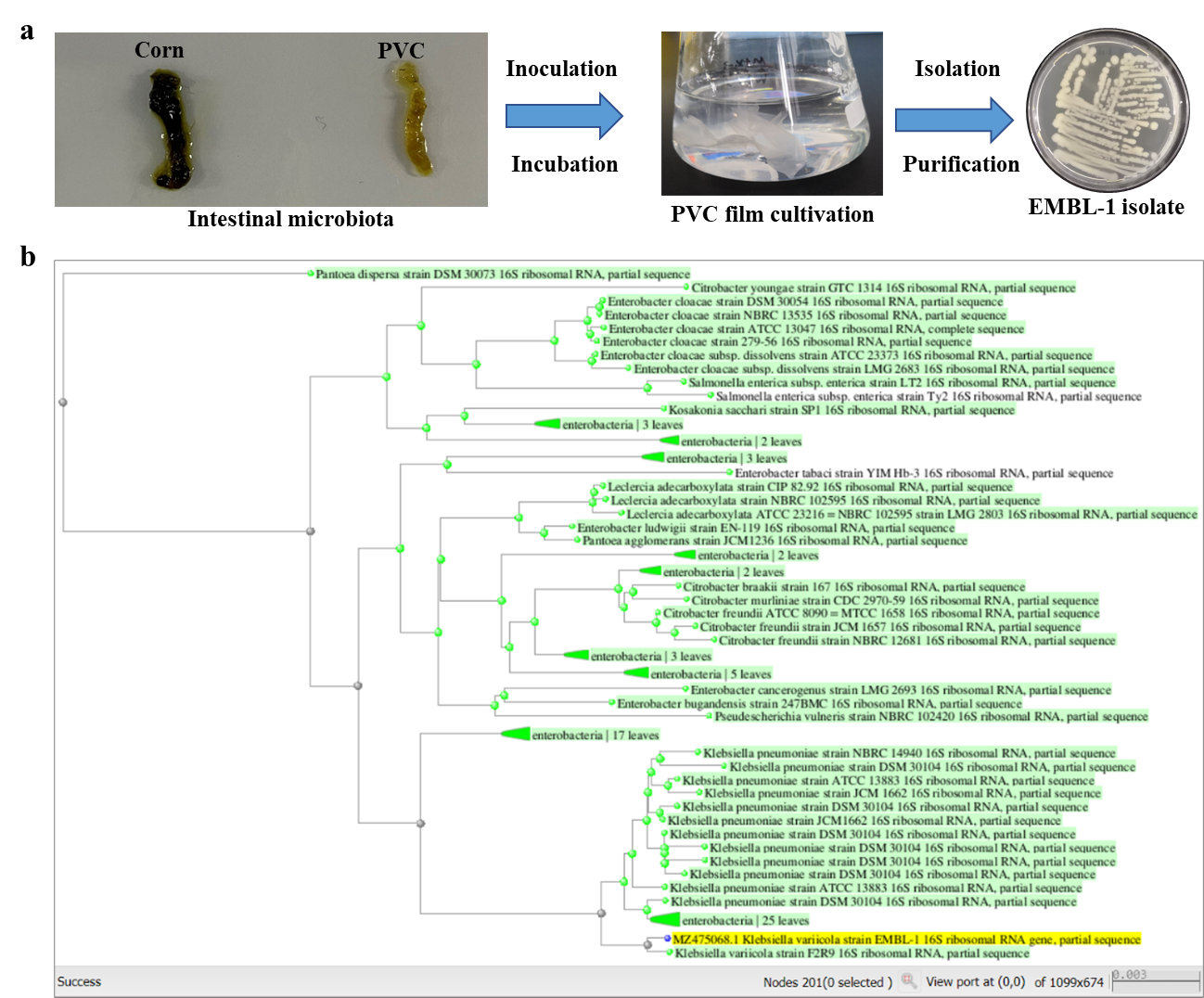

### Figure S3 The degradation results of strain EMBL-1 on PVC film

a, the SEM image of the cracks and pits formed on the PVC film by EMBL-1 strain on day 90. b, water contact angle test results of PVC film after co-culturing with strain EMBL-1.c, Tensile strength of PVC film in the control group and the EMBL-1 group after 90 days. d, the growth status of the EMBL-1 strain using SO as the substrate in 65 days. e-f, thermal gravimetric analysis (TGA) results of PVC film in control and EMBL-1 groups during 90 days.

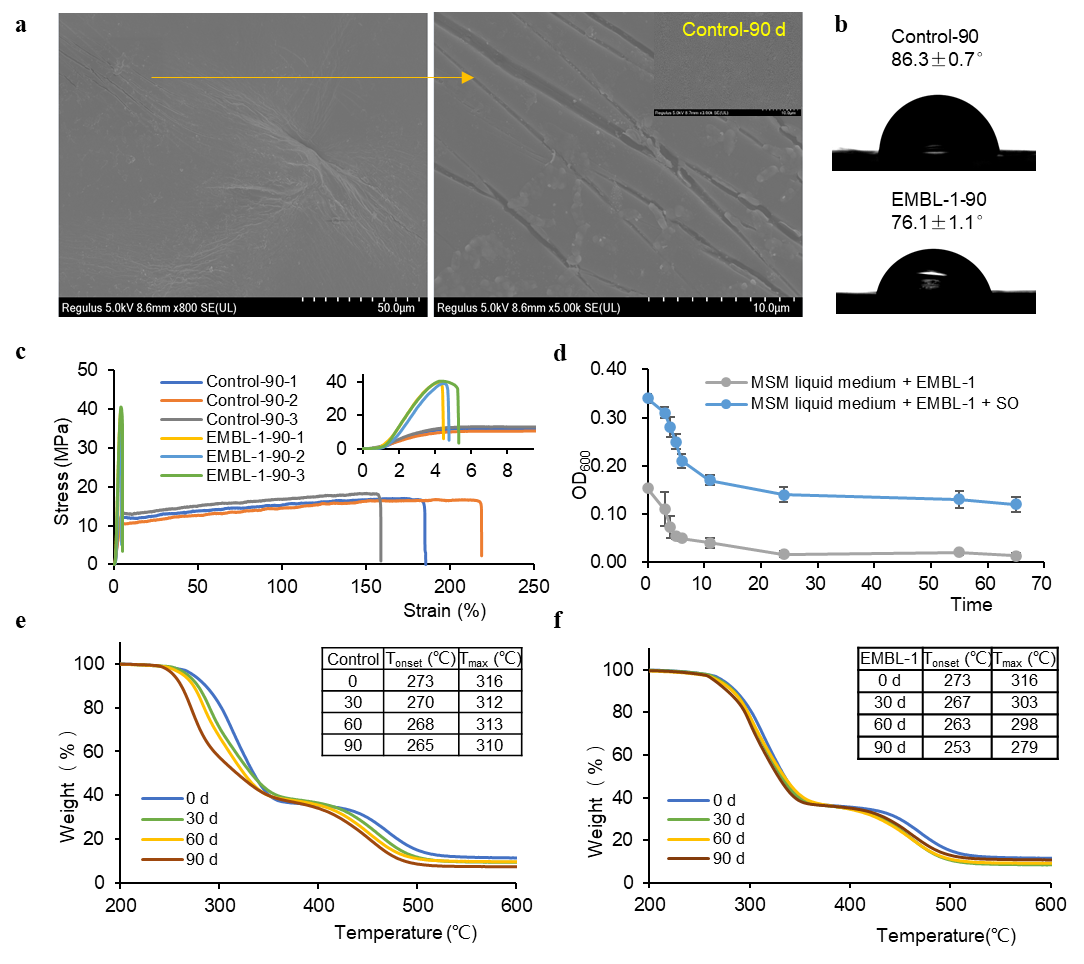

### Figure S4 The spectrum of NMR experiments of PVC

a, the DOSY Spectrum of pure PVC in control and EMBL-1 groups during 90 days. b, the 2D ^1^H-^1^H COSY spectrum of pure PVC in EMBL-1 groups during 90 days. c, the 2D ^1^H-^1^H NOESY spectrum of pure PVC in EMBL-1 groups during 90 days. d, the 2D ^1^H-^13^C HSQC spectrum of pure PVC in EMBL-1 groups during 90 days. e, the 2D ^1^H-^13^C HMBC spectrum of pure PVC in EMBL-1 groups during 90 days.

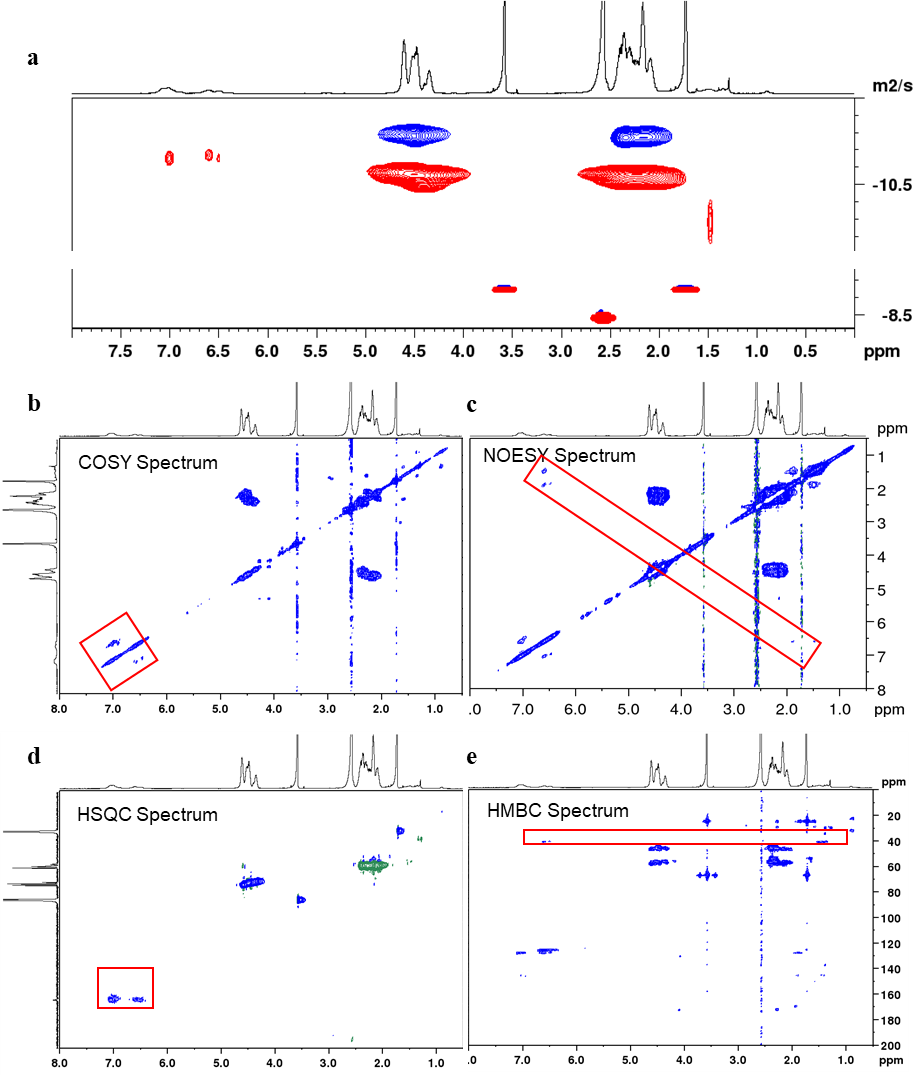

### Figure S5 Detection results of degradation products of PVC film by GC-MS

a, the TIC diagram of degradation products of PVC film in two groups by the time of 90 d. b-f, the mass spectrum and structural formula of the potential degradation products 1-5.

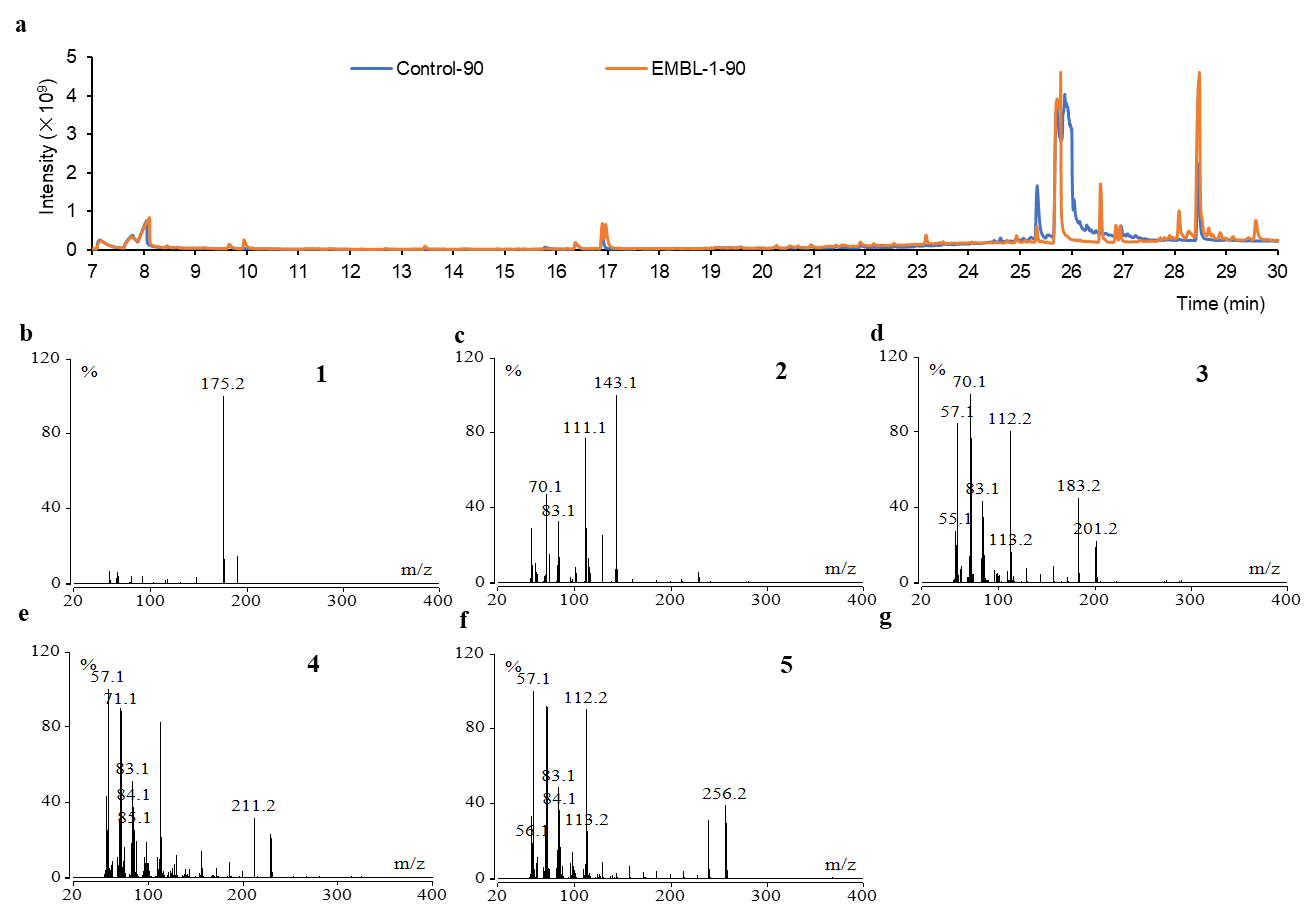

### Figure S6 The detection and identification results of additives in PVC film and the degradation activity of EMBL-1 strains on main additives

a, the Py/GC-MS diagram of three additives in PVC film. b, the TIC diagram of three additives identification in PVC film. c-e, the mass spectrum of three additives. f, the result of degradation activity of EMBL-1 strains on three additives.

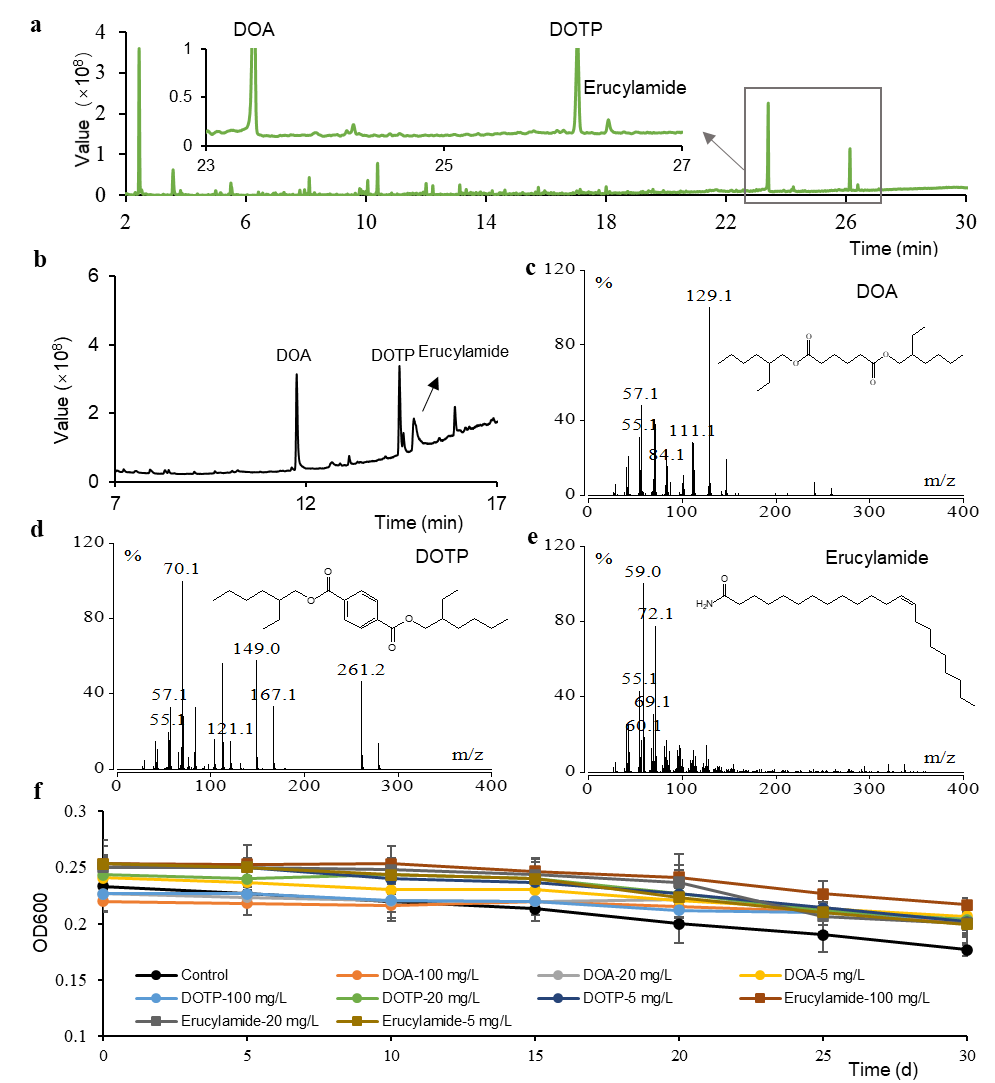

### Figure S7 The growth curve of the EMBL-1 strain and the concentration of extracted protein in the proteomic experiment

a, growth curve of strain EMBL-1 under different carbon source conditions. b, the concentration of proteins in four groups detected by Bradford method.

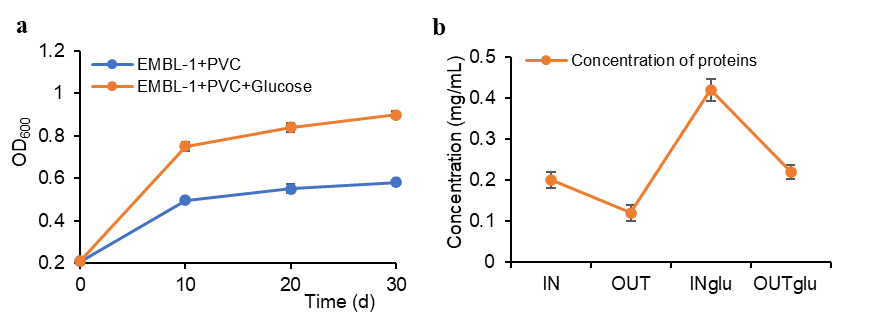
